# The Incoming Influenza Genome Assembles a Host RBP Network that Orchestrates Viral RNA Synthesis

**DOI:** 10.64898/2025.12.10.693272

**Authors:** Stephen Clarke, Deep B. Patel, Andrea Ascura, Caroline L. Wiser, Jennifer L. Pennise, Samantha M. Lisy, Manuel Ascano

## Abstract

Influenza A virus (IAV) replication initiates within minutes of entry, yet the earliest host determinants acting on the incoming viral genome remain undefined. Here, using VIR-CLASP to capture protein interactions with progenitor vRNA prior to any detectable viral transcription, we map the first host-viral RNA interface and identify ∼700 human RNA binding proteins (RBPs) proactively recruited by the negative-sense genome. These factors are enriched in pathways essential for RNA metabolism, chromatin remodeling, transcriptional regulation, and nuclear condensate organization, revealing that the vRNA engages a far broader host network than previously appreciated. Functional interrogation of top candidates demonstrates that GMPS, TOP2A, SRRM2, and SPEN serve as critical proviral factors operating at distinct stages of the IAV synthesis program: GMPS and SPEN are required for efficient production of vRNA, cRNA, and mRNA; TOP2A promotes mRNA capping and early transcript accumulation; and SRRM2, together with SR-proteins, coordinates splicing of viral M and NS transcripts. These findings support a model in which the incoming vRNA acts not merely as a transcriptional template but as a scaffold that initiates assembly of nuclear machinery required for productive viral replication. By defining the pre-replicative viral RNA interactome and its functional consequences, this work exposes an unrecognized layer of host control over IAV permissivity and establishes new points of vulnerability for antiviral intervention.

## INTRODUCTION

Influenza A virus (IAV) remains a major respiratory pathogen responsible for significant morbidity and mortality worldwide, causing seasonal epidemics and unpredictable pandemics. IAV infection is traditionally understood to begin with the binding of its hemagglutinin (HA) protein to sialic acid receptors on host cell surfaces^1^, a critical determinant of host tropism and viral transmission^2–10^. Specifically, human-adapted strains like H3N2 preferentially recognize α-2,6-linked sialic acids, while avian strains bind to α-2,3-linked receptors. However, mounting evidence suggests that cell surface receptor availability alone may not fully explain the complexities of viral permissivity and tissue tropism.

Disparities in replication competence are observed between different cell types^11,12^, suggesting that other factors including host-specific intracellular components influence the success of IAV replication. After viral entry, the viral ribonucleoprotein (vRNP) complex, containing the viral RNA (vRNA) genome, is released into the cytoplasm prior to transport into the nucleus wherein replication commences. The IAV life cycle hinges on the intricate and tightly regulated molecular synthesis of these three distinct RNA species. The stages of replication involve transcription of incoming vRNA genome into either complementary RNA (cRNA) or messenger RNA (mRNA), by the viral RNA-dependent RNA polymerase (vRdRp), serving as templates for progeny vRNA or viral protein synthesis, respectively. The transcription process itself requires both the vRdRp and interplay with host machinery which regulate viral replication. Such host factors play a role in facilitating replication and cRNP assembly^13^, capping of IAV mRNA through cap-snatching^14–16^, and alternative splicing of segments 7 (matrix protein; M)^17,18^ and 8 (nonstructural protein; NS)^19^ in a process entirely dependent on the host cell’s splicing machinery^20^. In a separate, primer-independent process, the same vRdRp complex shifts its function to replicate the incoming vRNA using the cRNA as template^21–23^. Therefore, the replication of IAV requires a combination of protein-protein and protein-RNA interactions, the former of which has been the major topic of study by others^24–42^.

Since viral RNA replication processes are dependent on viral and host proteins, we reasoned that, in the absence of phenotypical differences in cell surface receptor distribution that may occur between cell types or differences between strain invasion efficiency, the major driving force behind viral replication is the intracellular landscape the RNA genome encounters within the host. While numerous studies have examined the interplay between IAV and host proteins, very little is known about cellular proteins that directly interact with – and functionally influence – the incoming segmented genome of the virus. Studying the pre-replicative interactome of IAV may reveal key host factors – both known and novel – that act on the virus during this early, critical stage of its life cycle.

The initial interaction between the incoming viral RNA genome and the host’s cellular machinery, particularly host RNA-binding proteins (RBPs), represents a crucial, yet understudied, stage of the IAV life cycle and the role of host RBPs in regulating viral permissivity remains a significant knowledge gap. To address this, we employed VIR-CLASP (Viral Crosslinking And Solid-phase Purification) to discover the interactome of the incoming IAV genomic vRNA with human host proteins in lung-derived cells (A549). Our analysis uniquely identifies a suite of approximately 700 host proteins that interacts with the incoming viral genome prior to the onset of replication. This dataset reveals the specific host proteins that are "recruited" by the viral genome to prepare the cellular environment for successful replication. This includes factors involved in transcription regulation, RNA modification, splicing, and chromatin remodeling, such as TP53, FMR1, SRSF3, CHD1, and TOP2A, suggesting that IAV strategically subverts these host processes for its own benefit.

By mapping the cell-specific vRNA interactome, our study provides a novel perspective on IAV replication, moving beyond a simple template-driven process to one where the incoming viral genome is proactive in orchestrating the necessary cellular conditions. We show that the incoming vRNA acts as a scaffold, recruiting host factors essential for the synthesis of all intermediary viral RNA species (mRNA and cRNA). Our validation of specific host RBPs reveal pro-viral host factors like GMPS, TOP2A, SRRM2, and SPEN influencing viral replication at the RNA level. Their effects span across various stages of the IAV replication cycle, including initiation, cap-snatching, and splicing. Thus, the IAV genomic vRNA acts not only as template, but as the proactive scaffold for assembling the necessary components for the synthesis and replication of subsequent RNA necessary for production of progeny virus.

These findings reveal the nuanced and critical role of post-entry RBP interactions in regulating IAV replication and highlight the potential of these proteins as novel targets for antiviral intervention. Furthermore, host RBP functions in other viruses may be conserved and provide intriguing targets for clinical research. By elucidating these intricate host-virus interactions, our research reveals new strategies to disrupt the early stages of the IAV life cycle and limit this significant global health threat.

## RESULTS

### The negative strand of Influenza A Virus directly interacts with host factors prior to any viral transcription or replication event

To determine the host factors acting on the virus immediately after entry, we first aimed to identify a model system for studying the molecular basis of IAV replication. It is well established that IAV predominantly utilizes host sialic acid receptors^1,43–45^, amongst others^46^, for recruitment of virions to the cell membrane prior to entry and cytoplasmic release of vRNPs. Viral RNPs may be detected in the cytoplasm as early as several minutes post-binding to host cells and within the first hour of infection, are transported to the nucleus wherein viral replication starts by converting the incoming genomic vRNA (minus sense) to complimentary vRNA (cRNA) (plus sense) and viral mRNA (plus sense)^45^.

Due to the inherent differences in cellular proteomes and surface protein presentation, we evaluated the viral entry and replication in several cell lines, including A549, BEAS-2B, U2OS, Hep-2, and HuH-7 cells (data not shown). Using this data, we decided to monitor viral entry differences and establish equal entry conditions in A549 and BEAS-2B cells (Figure S1A). The difference in viral replication between the lung-derived A549 and BEAS-2B cells displayed a striking result – despite equal entry, viral RNA replication was notably impaired within BEAS-2B cells^47^. This effect was observed across all eight viral genome segments. Even with minimal RNA replication occurring within the BEAS-2B cells, no increase in titer was detected by TCID_50_ assay while A549 cells replicated IAV ∼1,000-fold relative to start (Figure 1A). We therefore focused on A549, given the demonstrated permissivity for H3N2.

**Figure 1.**
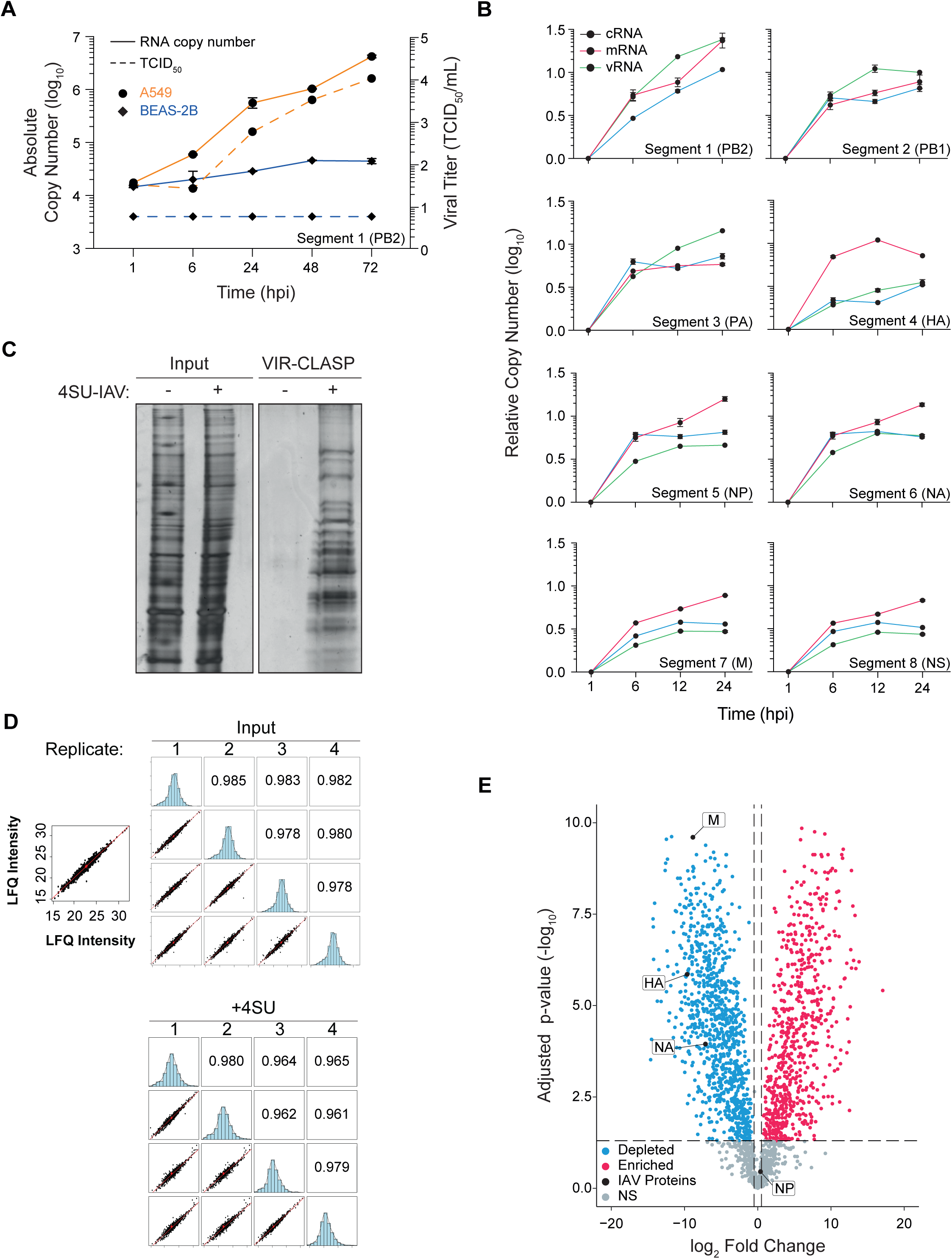
The pre-replicated Influenza A genome directly interacts with host proteins in human A549 cells. (A) IAV replication kinetics in entry-normalized human A549 (orange) and BEAS-2B (blue) cells over a 72 h period. Solid lines represent IAV segment 1 (PB2) absolute RNA copy number and dashed lines represent IAV titer as determined by TCID_50_ assay. (B) A549 cells were infected with IAV at MOI 0.1 and the RNA levels for each IAV segment and strand (c-, m-, and vRNA; blue, red, green, respectively) were monitored over a time-course of 24 h (1, 6, 12, and 24 hours post infection, hpi). All data normalized to the respective strand copy number at 1 hpi. (C) SDS-PAGE and silver stain of proteins captured by VIR-CLASP with 4SU-labelled IAV from A549 cells at 1 hpi. (D) Pearson correlation coefficients between all replicate samples of Input and +4SU in demonstrating reproducibility. Compare to correlation analysis showing differences between the identified proteins from Input, -4SU and +4SU samples in Figure S1. (E) Volcano plot analysis of IAV-A549 VIR-CLASP MS experiment showing enriched (red) or depleted (blue) proteins as compared to the input control. IAV proteins were also identified and indicated. N = 4. Error bars (in A and B) represent mean +/- SD, n = 3. Horizontal and vertical dashed lines (in E) indicate p-value 0.05 and Log_2_ Fold Change +/- 1 (2-fold), respectively.

We further delineated the viral RNA replication kinetics via strand-specific (viral v-, c-, and mRNA) quantification per segment over a 24 h period within the IAV-permissive A549 cell line – establishing a baseline for viral replication (Figure 1B). Studies have shown that vRNA replication does not correlate with cRNA^48–50^, which is suggestive of complex regulation of vRNA synthesis beyond just having template available. We demonstrated that all segments replicate vRNA in greater amounts than that of both c- and mRNA. NS1 yielded a significant spike in mRNA levels by 6 h post-infection^48–50^. Our observation in strand-specific replication kinetics of each segment reflect the observations by others using different IAV strains^50–52^. Segments encoding the vRdRp components produced more vRNA than c- and mRNA, while those encoding the viral HA, nucleoprotein (NP), and NS proteins produced more mRNA than v- and cRNA. The kinetics observations were mirrored in our data for A549 infected with H3N2 – indicating that the replication kinetics remain similar between these IAV strains. Taken together, our viral entry and replication results indicate that there are profound differences between A549 and BEAS-2B cells which dictate the outcome of IAV infection. Such differences extend beyond the potential differences in IAV receptor distributions as the infection differences persist when viral entry is equalized. Host intracellular factors such as RBPs may thus play a pivotal role in such cell-specific differences.

Using our viral and cellular model we applied VIR-CLASP^53,54^ on IAV-infected A549 cells. This method allows for unbiased isolation of RBPs directly interacting with incoming 4SU-labelled genomic vRNA. We previously established that IAV replication relies on progenitor vRNA prior to 3 hpi^54^ and, consistent with our previous study, other groups have also observed that the conversion of progenitor vRNA to cRNA and mRNA occur as early as 2 hpi^45,48,55^. For this reason, we infected A549 cells for 1 h prior to crosslinking and collecting the cells. The VIR-CLASP samples were processed and aliquots were fractionated by SDS-PAGE followed by silver stain (Figure 1C). The remainder of the recovered crosslinked RNP complexes were analyzed by LC-MS/MS and subsequent proteomics data analyses using the MSstats R package^56^.

To identify proteins interacting with the pre-replicative IAV genome, we performed proteomics analysis on the VIR-CLASP samples at 1 hpi. The raw proteomics data was processed by MaxQuant using a DDA pipeline with LFQ normalization applied. Both human and canine proteomes were supplied for the identifications to provide a stringent filter for removing any potential proteins that may originate from the host propagation line, MDCK. The IAV proteome for A/CA/07/04 (H3N2) was also supplied to account for any viral peptides detected in the samples as well as the sequence for Benzonase. This total protein identification data was processed using the MSstats R package to apply quantile normalization of the protein intensity values, imputation of missing values, removal of reverse/decoy peptides, exclusion of proteins identified by only 1 peptide. During this process PTXQC analysis was performed and used to qualify the integrity of the data (data not shown). Pearson correlation of the data revealed very high correlation (> 0.96) between sample replicates (Figure 1D). The correlation of the log2 intensity values of each sample (Figure S1C) demonstrated that the -4SU sample has low complexity as observed by the low correlation coefficient (0.364) with the input, the left-ward shift in the intensity distribution histogram and comprising 1,125 compared to 3,282 human proteins within the input. Low correlation was also observed in relation to the +4SU sample (2,275 proteins), indicating low similarity between -4SU and the input or +4SU samples. Conversely, the +4SU samples demonstrated higher complexity as observed in higher Pearson correlation with the input (0.607) and normal distribution of the intensity histogram. The left-ward shift in -4SU histograms indicates that our methodology has low-complexity, low abundance background. Together, this demonstrates that the ideal samples for comparison are the +4SU and input as the -4SU sample would decrease confidence in the validity of enrichment data.

The proteomics result show that the major IAV capsid proteins are depleted. Most notably, the HA and neuraminidase (NA) proteins (although not actively engaged with the viral genome) show that no significant amount of intact virus was present in the samples. Depletion of the M and NP proteins demonstrate that lysosomal fusion and deployment of the viral genome had occurred and vRNA decoupling from NP indicate that the vRNA is preparing to engage other factors for RNA replication^45,46,57^. We did not observe any detectable amounts of the viral RNA polymerase subunits (Polymerase Basic 2; PB2, Polymerase Basic 1; PB1, and Polymerase Acidic; PA proteins) nor its peptides within the raw mass spectrometry data. We identified ∼800 human proteins of which 676 were significantly enriched (adjusted p-value <= 0.05 corresponding to an FDR of 0.01% and a log2FC > 1) including several known host factors involved in IAV infection such as IFI16^54,58–60^, hnRNP proteins^58,61^, and NONO^61^ (Table S1).

### Host factors interacting with the IAV genome are from key cellular pathways exploited by IAV

We characterized the IAV VIR-CLASP interactome based on previous studies on IAV-host interactions. Our IAV interactome had 128 proteins (19%) overlapping with the virome of *Orthomyxoviridae*^62^ and 548 proteins (81%) unique to our dataset (Figure 2A, top). When compared to the well-studied host-IAV protein-protein interactome (PPI) (compiled from 17 curated studies^25–42^ by Chua *et al*., 2022), our data provides 521 (77%) unique proteins over the 155 proteins (23%) overlap (Figure 2A, bottom). Most of the shared proteins had previously reported protein-protein interactions with vRdRp PA^28,32,34,36,40^ protein and viral structural proteins (NP^29,36,40^ and NS1^26–32,34,36–39,42^). The IAV-host PPI was generated from studies using various methodologies, ranging from bioinformatics to immunoprecipitation and RNAi screens. Our study, specifically enriching for RNA-bound host proteins, expands on these previous studies by validating that the 155 shared proteins are RBPs (with potential protein-protein interactions) and the remainder are likely protein-binding alone.

**Figure 2.**
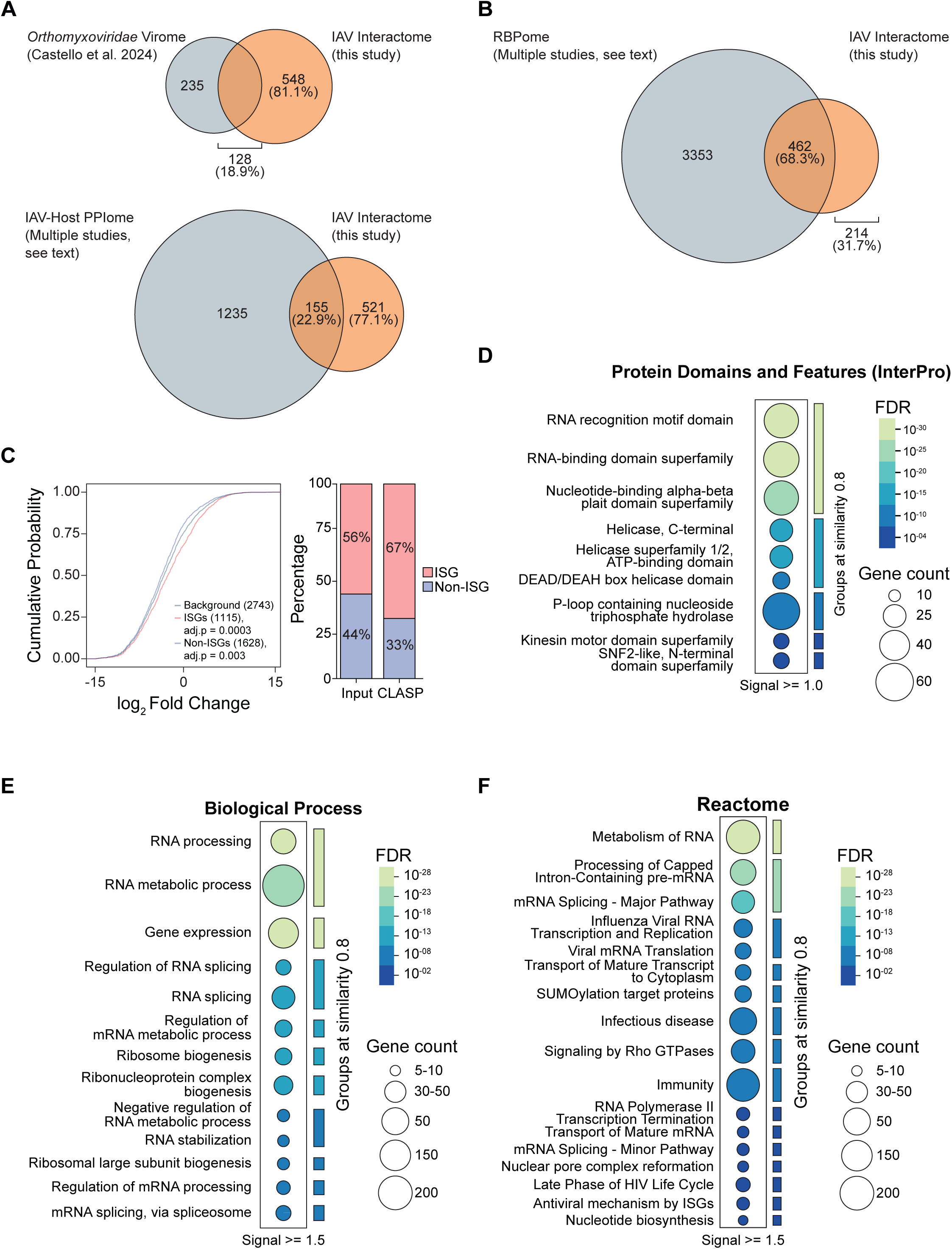
IAV genome recruits host RBPs from key RNA metabolic pathways. (A) Euler plot between IAV VIR-CLASP and *Orthomyxoviridae* interactome^62^. (B) Euler plot between this study’s IAV interactome and the RBPome as determined from previous studies^63,69^ (see text). (C) The incoming IAV genome enriches for proteins defined as ISGs, as compared to A549 cell lysate (input). Cumulative distribution analysis demonstrates enrichment of ISGs (adjusted p-value = 0.0003). The proportion of A549 input and VIR-CLASP interactome comprising of proteins identified as ISGs are displayed in accompanying bar plot. (D) InterPro analysis detailing protein binding motifs enriched within the IAV interactome. (E and F) Gene ontological analysis of IAV interactome based on Biological process annotations (E) and Reactome pathways (F). FDR = False Discovery Rate (in D, E, and F).

We further demonstrate that the majority of our interactome comprises of known RBPs when compared to the RBP2GO^63^ and RBPBase^64–69^ *Homo sapiens* RNA-Binding Proteome (RBPome) databases (Figure 2B). The IAV interactome comprised of 462 (68%) proteins annotated as RBPs while 214 (32%) proteins represent potentially novel RBPs. The distribution of RBPs was further characterized by Pfam identifiers associated with validated RNA binding domains (RBDs) present in RBPs (canonical RBD)^69^ and those not fully validated as RBDs conferring RNA binding properties to proteins (non-canonical RBD)^69–72^. The canonical RBDs such as RNA-recognition motif (RRM)^69,73,74^ and the K-homology domain^69,73–76^ have been well characterized as conferring RNA binding activity. It has also been demonstrated in recent years that non-canonical RBDs, such as sterile alpha motif^74^ and intrinsically disordered regions (IDRs) are also enriched within RBP candidates^70,77^. For this study, we classified proteins as “conventional”-RBPs (cRBPs) or “unconventional”-RBPs (ucRBPs) if they contained at least one canonical RBD or only the so-called non-canonical RBDs, respectively. Proteins not containing either grouping of domains or motifs were classified as ‘potential”-RBPs (pRBPs). We determined that the human RBPome (as defined in this study) comprises 83% pRBPs, 11% ucRBPs, and 6% cRBPs – in line with the RBP2GO characterization of the human proteome RBP distribution^72^. This distribution was preserved within the IAV interactome (82%, 9%, and 9%, respectively), with a slight increase in cRBP content (Figure S2A). The IAV interactome proteins shared with the RBPome comprised of 13% cRBPs, 11% ucRBPs, and 76% pRBPs while the potentially novel RBP population (214 proteins) had a distribution of only ucRBPs and pRBPs, 3%, and 97%, respectively. We further characterized the percentage of IAV interactome proteins annotated as interferon-stimulated genes (ISGs)^58^. Because ISG transcript stability is often regulated by RBPs, we investigated the pre-replicative IAV genome engagement with ISG proteins. We queried the Interferome database^78^ with our A549 input and IAV interactome, characterizing 56% and 67% as ISGs, respectively (Figure 2C). Cumulative Distribution Factor (CDF) analysis showed that enriched proteins are significantly represented by ISG proteins (adjusted p-value = 0.0003). The incoming vRNA therefore engages with predominantly cRBPs of which ISG proteins are enriched. Since we posit that RBPs engaging with the incoming vRNA regulate IAV infection and that ISGs coordinate viral infection responses, we selected this subset of the IAV VIR-CLASP interactome for further analyses (Supplemental Table 3).

We used the filtered IAV interactome to assess the proteins by Gene Ontology (GO) and Reactome pathway analysis^79–81^ using the STRING functional protein association analysis^82^. When assessing the data outputs from GO and Reactome pathway analyses, we looked at both the “strength” (proportion of term present in the IAV interactome compared to computational expectation) and “signal” (weighted harmonic mean between term “strength” and FDR). The signal metric aims to account for the potential of very large GO terms from masking the enrichment of terms with few members^82^. We found that most biological processes involved RNA metabolic processes (GO:0016070; GO:0006396), gene expression (GO:0010467), regulation of RNA splicing (GO:0043484), and, interestingly, regulation of topoisomerase activity (GO:2000373). Molecular function process enrichment included general RNA binding activity (GO:0003723), “ATP-dependent chromatin remodeler activity” (GO:0140658) and “poly(G) binding” (GO:0034046) while Reactome pathway enrichment included “processing of capped intron-containing pre-mRNA” (HSA-72203) splicing (HSA-72172, HSA-72163), RNA pol-II transcription termination (HSA-73856), SUMOylation (HSA-3108232, HSA-4570464), and “Influenza viral RNA transcription and translation” (HSA-168273) – the latter as expected for a permissive cell line such as A549.

We compiled all the information from the CDF, GO, and Reactome pathway analyses and summarized it into a more digestible network (Figure 3A and Table S5). The proteins were grouped into umbrella terms (large colored spheres) including “RNA Metabolism”, “Gene Expression”, “Regulation of Gene Expression”, “Immune System”, and “Infectious Disease”. Within each umbrella term we grouped GO terms and linked the network via shared genes (lines). This summarized network was used to select proteins representing each process for further validation, spanning log2FC values from 1.1 (1.2-fold) to 14 (196-fold), including RAN, GMPS, TOP2A, UBE2L3, WDR33, YTHDF2, SRRM2, SPEN, PARP4, IK, TP53, CHD1, TTF2, RBBP6, ATRX, IFI16, ZC3H13, and DDX56.

**Figure 3.**
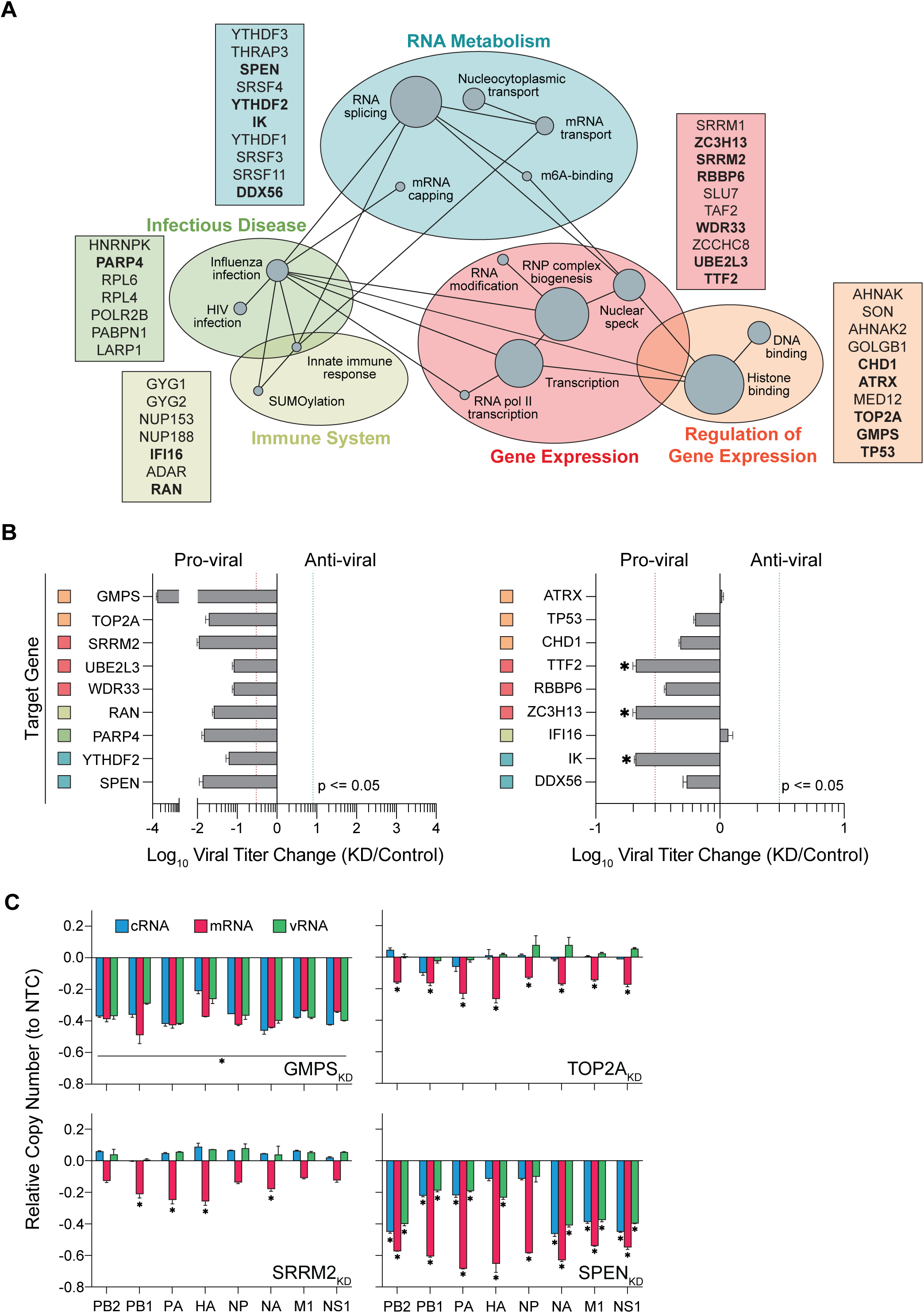
Candidate host proteins impact replication of IAV c-, m-, and vRNA. (A) Pathways of IAV interactome. STRING protein-protein interaction network analysis of IAV VIR-CLASP interactome filtered for known RBPs and ISGs. Proteins are grouped by pathway and functional roles (large spheres) with each node (grey circle) denoting a GO term within the grouping. Proteins listed in boxes represent members within each grouping. In bold proteins selected for further functional validation. (B) siRNA knockdown of candidate proteins within A549 cells for three days were subjected to IAV infection and IAV titer measured by TCID_50_ at 24 hpi. Titer was determined by collection of culture medium from infected or mock infected cells under each transfection condition and performing TCID_50_ assay using MDCK cells. TCID_50_ results are presented relative to siRNA control infected with IAV. (C) Relative strand-specific viral RNA copy number upon target knockdown to control (non-targeting siRNA treated) infection at 24 hpi. Bars represent cRNA (blue), mRNA (red) and vRNA (green). Error bars represent SD, n = 3, * = p <= 0.05 (in B and C). Dotted lines represent a 10-fold difference from control (in B).

### VIR-CLASP candidates are important for infectious IAV virion production

The selected host proteins were screened for effects on IAV infection. Gene knockdown of each target was performed followed by infection with IAV for 24 h prior to viral titer determination by TCID_50_ (Figure 3B). ATRX and IFI16 resulted in an increase in titer (∼1.3-fold) indicating these host factors normally restrict IAV replication. IFI16 acts as an anti-viral restriction factor as previously demonstrated^59,60^, however the knockdown did not increase the IAV titer significantly. This may be due to the already permissive nature of A549 cells wherein the expression of IFI16 mRNA is low (22-fold lower) compared to an immune-activated, IAV-restrictive cell line such as BEAS-2B as determined by RT-qPCR (data not shown). Conversely, the known pro-viral proteins, CHD1^83–85^ and RAN^86,87^, decreased the titer as expected. CHD1 is involved in the degradation of RNA Polymerase II (Pol II) and subsequently may act to reduce the anti-viral response^83^. Additionally it is suggested that CHD1 may be involved in recruiting the viral polymerase to positively modulate its activity and, despite IAV not incorporating into the host genome, chromatin remodeling factors such as CHD1 and CHD6^88,89^ may play a role in engaging the replicating viral genome with host gene expression machinery necessary for viral RNA replication. Our IAV interactome enriched several host proteins involved in chromatin remodeling, mirroring the notion that this pathway is important for IAV replication. Our screening most notably showed that knockdown of TOP2A, SRRM2, and SPEN resulted in ∼100-fold and GMPS ∼10,000-fold decrease in viral titer, presenting intriguing targets for further study within the context of IAV replication. All siRNA treatments had no effect on cell viability (data not shown) while displaying at least significant reduction in target gene mRNA (Supplemental Figure S3A).

TOP2A, while canonically a DNA topology enzyme^90^, significantly impacts RNA-related processes, particularly transcription through its relationship with RNA Pol II^91,92^. Given IAV’s nuclear replication cycle and its reliance on host RNA Pol II for viral transcription^15,93–95^, TOP2A’s function in maintaining a permissive chromatin environment may be co-opted by the virus.

The nuclear Serine/Arginine Repetitive Matrix 2 (SRRM2) protein is primarily involved in RNA processing, particularly mRNA splicing and alternative splicing^96^. As a core component of the spliceosome, SRRM2 recognizes splicing elements on precursor RNA and organizes dynamic splicing condensates, or nuclear speckles, through liquid-liquid phase separation driven by its IDRs and RNA interactions^96–98^. Given that IAV lacks its own splicing machinery and is entirely dependent on usurping the host spliceosome to perform alternative splicing of segment 7 (M) and segment 8 (NS), both essential for the viral life cycle^55^, SRRM2, therefore may act as a core scaffold protein highly likely to be directly co-opted by IAV to facilitate efficient viral mRNA processing and protein production as shown by our VIR-CLASP data.

Split End homolog (SPEN) functions as a large, highly conserved RNA-binding transcription repressor, with its activity heavily reliant on interactions with various RNA molecules. SPEN coordinates rRNA synthesis in endothelial cells by repressing noncoding promoter RNA, a long non-coding RNA, which is crucial for ribosome biogenesis and angiogenesis^99^. This function is indirectly but essentially critical for IAV replication, as the virus is entirely dependent on the host cell’s ribosomes for the massive synthesis of its viral proteins^55^. Furthermore, SPEN possesses RRMs and is involved in silencing endogenous retroviruses by binding to viral RNA and recruiting chromatin silencing machinery, highlighting its broad impact on RNA-mediated gene regulation and genomic defense^100^.

Guanosine Monophosphate Synthetase (GMPS) is a crucial enzyme in *de novo* purine metabolism^101,102^. The *de novo* catabolism of purine nucleotides has been heavily implicated in IAV replication as demonstrated through the inhibition of IMP dehydrogenase (upstream of GMPS) by pharmacological agents such as ribavirin. GMPS has recently been demonstrated to also have RNA-binding activity via pCLAP^103^, RICK^104^, OOPS^105^, XRNAX^106^, PTex^107^, and eRIC^108^.

### Knockdown of candidate host factors impact replication of IAV vRNA, cRNA, and mRNA

To further assess the role of the top hits (GMPS, TOP2A, SRRM2, and SPEN) on IAV replication, we performed segment-specific and strand-specific RT-qPCR (ssRT-qPCR) – evaluating the effects of each host candidate protein on IAV v-, c-, and mRNA (Figure 3B and Figure S4A). We initially studied the effects on each RNA strand at 24 hpi for several of the candidate proteins from the knockdown screening assay. Not all segments or RNA species were affected in the same way (Figure 3B). Most notably, GMPS and SPEN reduced all strands for all eight segments (although SPEN affected viral mRNA more) while TOP2A, and SRRM2 predominantly reduced viral mRNA relative to the siRNA control. The candidate proteins affect different stages of the viral replication process, impacting one or all RNA species. It must be noted that any impact on viral cRNA production would also impact vRNA as the production of new vRNA is contingent on cRNA template. Significant impacts on viral mRNA may have the same effect if viral protein synthesis is reduced significantly due to their involvement in downstream c- and vRNA synthesis. Therefore, a 24 hpi snapshot of viral replication may mask such effects. We aimed to unveil said effects by monitoring viral c-, m, and vRNA replication at earlier time-points.

### GMPS is crucial for early steps in viral RNA synthesis

To study the effects of GMPS knockdown on IAV replication at earlier times (prior to 24 hpi) we monitored viral RNA replication kinetics at 1, 6, and 12 hpi (Figure 4A). We found that depletion of GMPS reduced the copy number of each RNA strand and, while normally by 6 hpi significant levels of v-, c-, and mRNA are detectable, with GMPS knockdown only by 12 hpi were RNA species detected as the rate of replication was reduced (Figure S4B). This indicates that GMPS is necessary for efficient replication of IAV from the progenitor genome prior to protein synthesis. The early effects on viral replication likely points to the role guanosine plays in the priming of viral RNA essential for initiation of viral RNA transcription events^55^.

**Figure 4.**
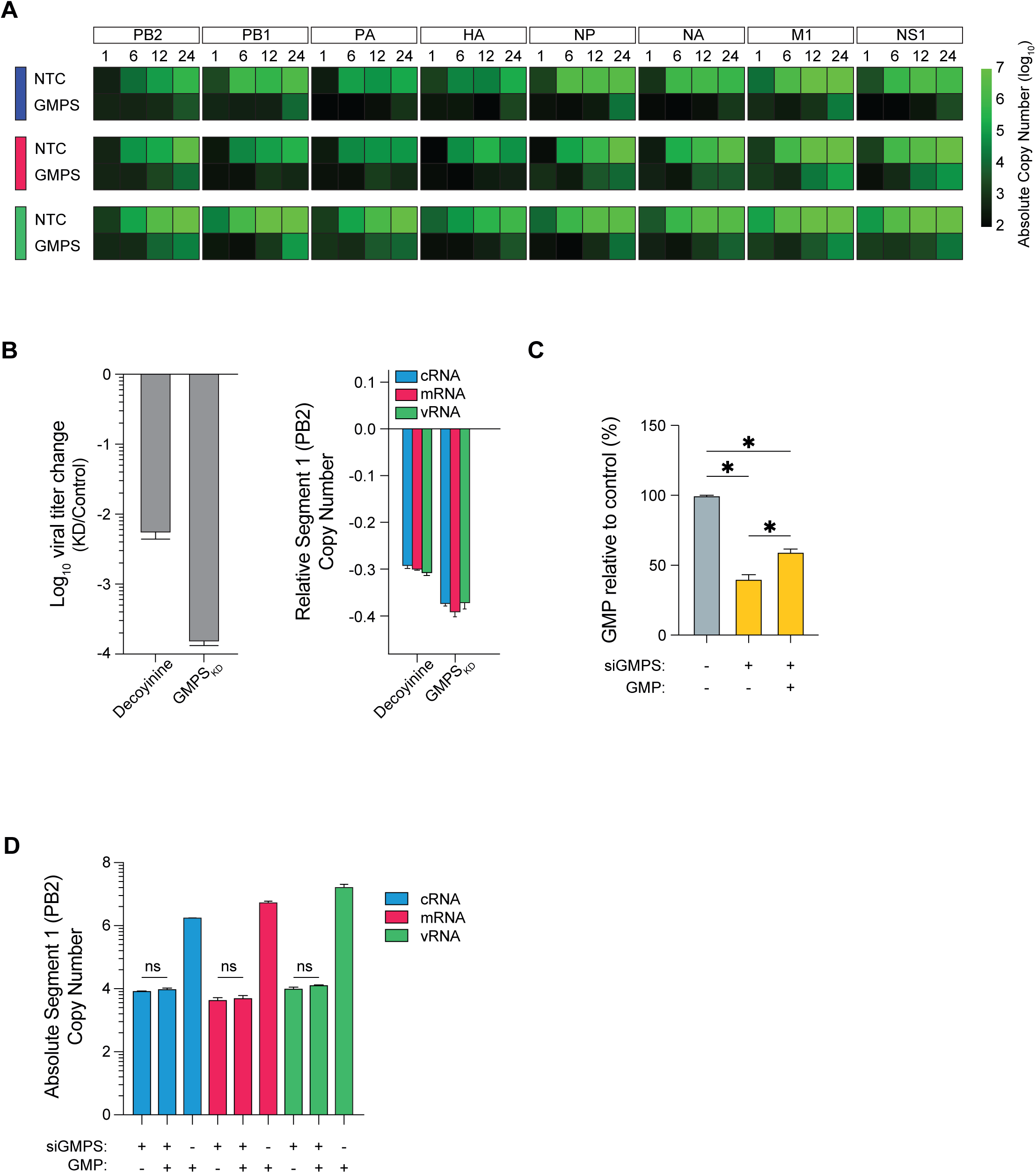
GMPS catalytic activity is required for IAV replication. (A) Heatmap of IAV replication kinetics under wild-type and GMPS knockdown conditions were monitored for each segment’s various RNA strands over a 24 h period (1, 6, 12, and 24 h). RNA absolute copy number is represented from low (black) to high (bright green). (B) The effect of GMPS catalytic activity was assessed by addition of the GMPS inhibitor decoyinine (at IC_90_ = 1.79 µM) to A549 cells prior to infection with IAV. The effect on IAV titer was measured by TCID_50_ assay (left) and the effect on RNA replication by ssRT-qPCR for viral segment 1 (PB2) (right). (C) Relative intracellular GMP levels as compared to untreated control cells. GMP was extracted from A549 cells treated with GMPS siRNA for 72 h. Exogenous GMP was supplied 16 h prior to collection followed by extraction of free nucleotide-phosphates and assessment by RP-HPLC. (D) Viral RNA replication under GMPS knockdown was monitored upon exogenous GMP supplementation (200 µM). Error bars represent mean +/- SD, n = 3. * = p <= 0.05 (in B and C)

Knowing that GMPS is essential for *de novo* biosynthesis of GMP^101,102^, providing the substrate for GTP synthesis, we wanted to study whether our knockdown of GMPS and the concomitant reduction in cellular GMP/GTP pool has a generalized negative impact on RNA synthesis. To do so, we first sought to compare GMPS knockdown to pharmacological inhibition of GMPS, thus leaving the protein levels intact while catalytic activity is impeded. GMPS activity may be inhibited by the adenosine analog, decoyinine^102^. We demonstrate that decoyinine has no effect on cell viability (see Figure S4B) while reducing viral segment 1 (PB2) (as proxy for viral RNA replication) c-, m-, and vRNA levels and titer (∼100-fold) (Figure 4B). This is intriguing since purine biosynthesis occurring via the salvage pathway can maintain intracellular GMP levels in absence of *de novo* pathway, provided there is a source of purine bases (adenine and guanine) ^109,110^.

IAV replication hinges upon the incorporation of a pppApG dinucleotide primer, from cellular GTP and ATP, which initiates replication of vRNA and cRNA^111,112^. Given that GMPS engages with the IAV vRNA and GMP is required for viral replication, we were interested in knowing whether GMP synthesis needed to occur during engagement with vRNA. We show that GMPS knockdown for 72 h reduced the intracellular GMP levels which we can partially recover by supplementation of exogenous GMP (Figure 4C). The replication of IAV was monitored by ssRT-qPCR within the knockdown and GMP-supplemented conditions which show that GMP supplementation has no effect within the knockdown (Figure 4D). Thus, our data show that impairing GMP synthesis by GMPS depletion or inhibition severely compromises IAV replication likely through disrupting replication initiation. Yet restoring total cellular GMP levels to near basal levels did not rescue IAV replication in GMPS-depleted cells.

### TOP2A is required for viral mRNA transcription and capping

Next, we assessed the effects of TOP2A depletion on viral RNA synthesis during the first 24 h. As with GMPS, we performed a replication kinetics assay (Figure 5A). We observed an early decrease in cRNA and vRNA, which recovered by 24 hpi while the most distinct effect was the specific reduction in viral mRNA.

**Figure 5.**
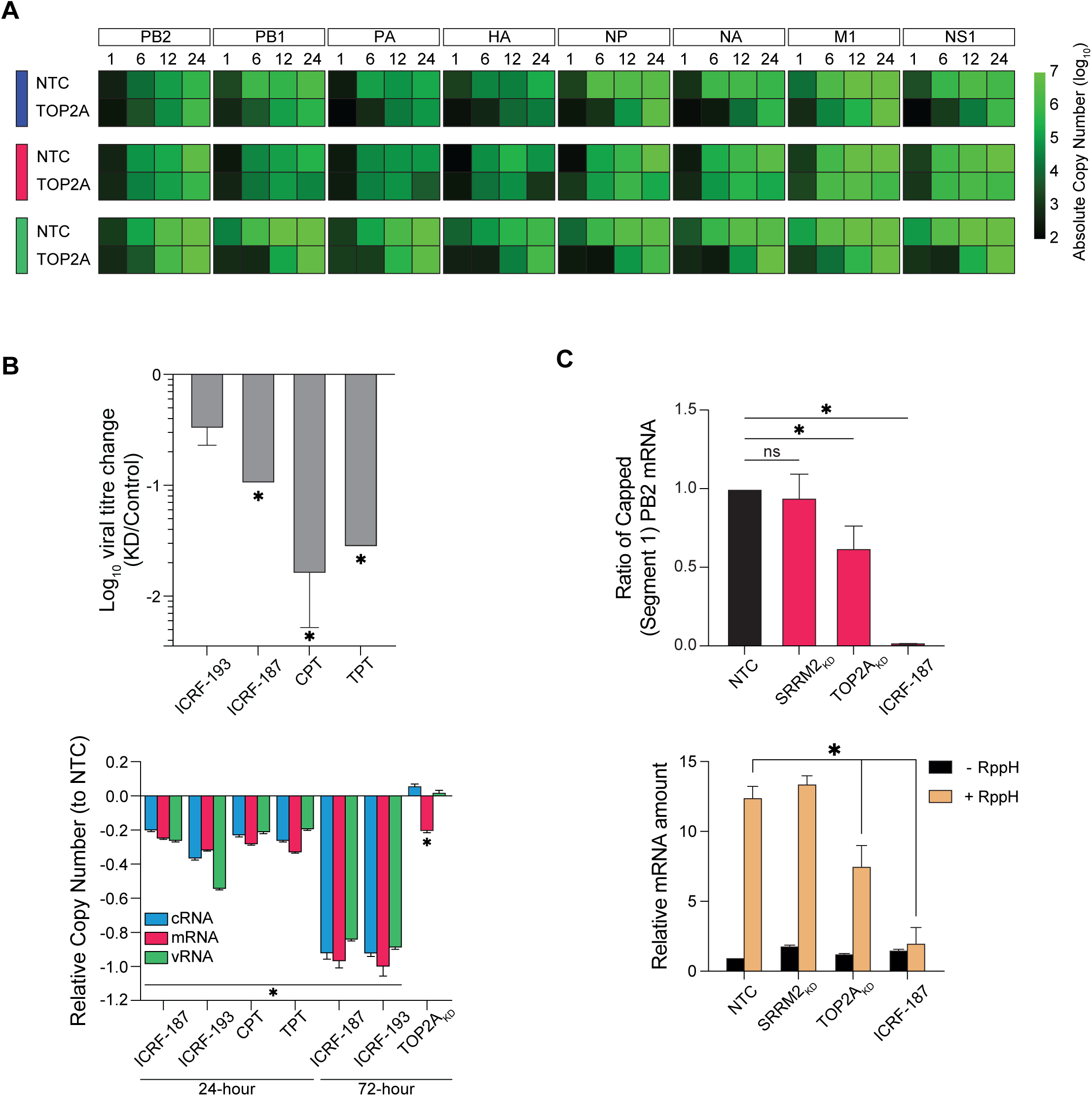
TOP2A is required for effective capping of viral mRNA. (A) Heatmap of IAV replication kinetics under wild-type and TOP2A knockdown conditions were monitored for each segment’s various RNA strands over a 24 h period (1, 6, 12, and 24 h). RNA absolute copy number is represented from low (black) to high (bright green). (B) The effect of TOP2A catalytic activity was assessed by addition of the topoisomerase inhibitors ICRF-193, ICRF-187, CPT, or TPT to A549 cells prior to infection with IAV (24 h or 72 h). The effect on IAV titer was measured by TCID_50_ assay (top) and the effect on RNA replication by ssRT-qPCR for IAV segment 1 (PB2) (bottom). (C) TOP2A influences capping of viral mRNA as demonstrated by cap-interference RT-qPCR. Data represents the ratio between uncapped viral mRNA within RppH-treated and untreated samples. The relative viral mRNA within each sample (bottom) indicates the necessity of RppH treatment for uncapped mRNA detection. Data was normalized to total viral mRNA in each sample prior to ratio calculations. Error bars represent mean +/- SD, n = 3. * = p <= 0.05 (in B, C, and D)

TOP2A significantly impacts RNA-related processes, particularly transcription^92^ and promote productive transcription elongation by facilitating the release of paused RNA Polymerase II (Pol II)^91^. TOP2A further exhibits RNA binding activity^65,67,103,105–107,113–116^ and participates in functions involving RNA-dependent proteins. Given IAV’s nuclear replication and its reliance on host RNA Pol II for viral transcription^15,93–95^, TOP2A’s function in maintaining a permissive chromatin environment may be co-opted by the virus.

To understand the role of TOP2A in IAV replication via engagement of the incoming vRNA we first investigated whether TOP2A catalytic activity influences IAV replication. We treated cells with topoisomerase inhibitors (ICRF-187, ICRF-193, camptothecin (CPT) and topotecan (TPT)) concomitantly with IAV infection for 24 h or 72 h, prior to infection with IAV for 24 h to compare the consequences of catalytic inhibition versus depleting TOP2A (Figure 5B). Viral titer was determined by TCID_50_ assay at 24 hpi and showed that all inhibitors reduced titer (top panel). Viral strand-specific RNA levels for segment 1 (PB2) (bottom panel) showed that all three strands (c-, m-, and vRNA) are significantly reduced in the 24 h and 72 h inhibitor treatments. ICRF-187 and the more potent form ICRF-193 both showed time-dependent reduction in viral RNA with the 72 h treatment rendering the RNA species near zero. However, the CPT and TPT 72 h treatments resulted in severe loss of cell viability (data not shown). Both ICRF-187 and ICRF-193 inhibit TOP2A/B activity traditionally through locking the enzyme in place on the DNA template preventing its release^92^. Since TOP2A was found to interact with the incoming IAV genome, simple depletion of TOP2A could result in a different outcome from use of catalytic inhibitor treatment – as they would lock TOP2A in place on the vRNA in a so-called rigor complex, preventing vRdRp progression during c- and mRNA synthesis. This may explain why all three RNA species are reduced as opposed only mRNA in the TOP2A knockdown, as the absence of TOP2A protein should not interfere with vRdRp-catalyzed viral cRNA synthesis. The formation of the rigor complex on the vRNA further emphasizes the direct interaction of TOP2A with vRNA.

The IAV vRdRp has been demonstrated to interact with the C-terminal domain (CTD) of RNA Pol II and this interaction is suggested to facilitate cap-snatching of host nascent mRNA 5’-caps^15,16^. It has also been demonstrated that TOP2A may interact with RNA Pol II and result in Pol II stalling^117^. These previous findings combined with our observation that TOP2A directly interacts with the IAV vRNA suggests a potential model wherein an intricate interplay between the IAV vRNA, vRdRp, TOP2A, and RNA Pol II may facilitate cap-snatching.

We further assessed whether TOP2A may be involved in viral cap-snatching by developing an assay to measure the level of capped viral transcripts. First, we isolated mRNA by oligo-dT pulldown followed by 5’-adapter ligation to samples either treated with or without RppH. The RppH would release 5’ cap moieties from mRNA, leaving a 5’-pN available for splint-aided adapter ligation. Compared to other approaches in measuring mRNA capping^118–133^, this approach first enriches mRNA, provides enhanced specificity and efficiency (splint hybridization aligns adapter to target), the DNA splint may be removed by DNase digestion – preventing non-specific amplification during qPCR, no secondary structure interference, and provides quantitative sensitivity via qPCR sequence-specific analysis of uncapped RNA targets. Following ligation, the samples are reverse transcribed, and cDNA quantified using target- and adapter-specific primers by qPCR. The difference in cDNA levels between RppH-treated and -untreated samples (normalized to total viral RNA) provides the quantity of originally capped mRNA^134^. We demonstrate that TOP2A knockdown reduced the level of capped viral mRNA by approximately 50% versus control while SRRM2 (a host factor involved in the downstream processing of mRNA via splicing) knockdown had no effect (Figure 5C). Interestingly, inhibition of TOP2A activity by 72 h treatment with ICRF-187 completely ablated capped viral mRNA highlighting that the inhibitor-induced formation of TOP2A rigor complexes on target DNA/RNA prevents production of *all* viral RNA species – distinct from TOP2A depletion which reduces only viral mRNA synthesis and capping.

### Splicing of IAV mRNA is directed by SR-proteins and SPEN

Serine/Arginine Repetitive Matrix 2 (SRRM2) is a nuclear protein primarily involved in RNA processing, particularly mRNA splicing and alternative splicing^96^ – which are required host processes for IAV replication^20^. Whereas Split End homolog (SPEN) functions as a large, highly conserved RNA-binding transcription regulator, with its activity heavily reliant on interactions with various RNA molecules, and has been demonstrated to interact with RNAs like X-inactive specific transcript (*Xist*) RNA^135^. Recent work suggests that the Repeat A domain in *Xist*, although critical for SPEN recruitment, does not directly interact with SPEN but rather SR-proteins. It has been shown that SR-proteins, such as SRSF1, directly interacts with SPEN and aid in its recruitment to *Xist* Repeat A domain^136^. Given that IAV requires host splicing machinery^20,45,50^ we next wanted to study the replication kinetics under SRRM2 and SPEN knockdown (Figure 6A). We show that SRRM2 knockdown reduced viral mRNA levels of segment 2-4 predominantly, however c- and vRNA were also reduced at earlier time-points for all segments before recovery by 24 hpi. Despite the reduction in RNA replication, the ∼20% reduction is likely not enough to overcome the overall replication of IAV and instead results in a milder infection which eventually recovers. The rate of replication (Figure S6A) is reduced compared to control up to 6 hpi, whereafter it increases to that of control. We also observed similar trends within the SPEN knockdown where replication of all viral RNA species was significantly impaired and the absolute copy number of c-, m-, and vRNA remain ∼1,000-10,000-fold lower than that of control – similar to GMPS knockdown.

**Figure 6.**
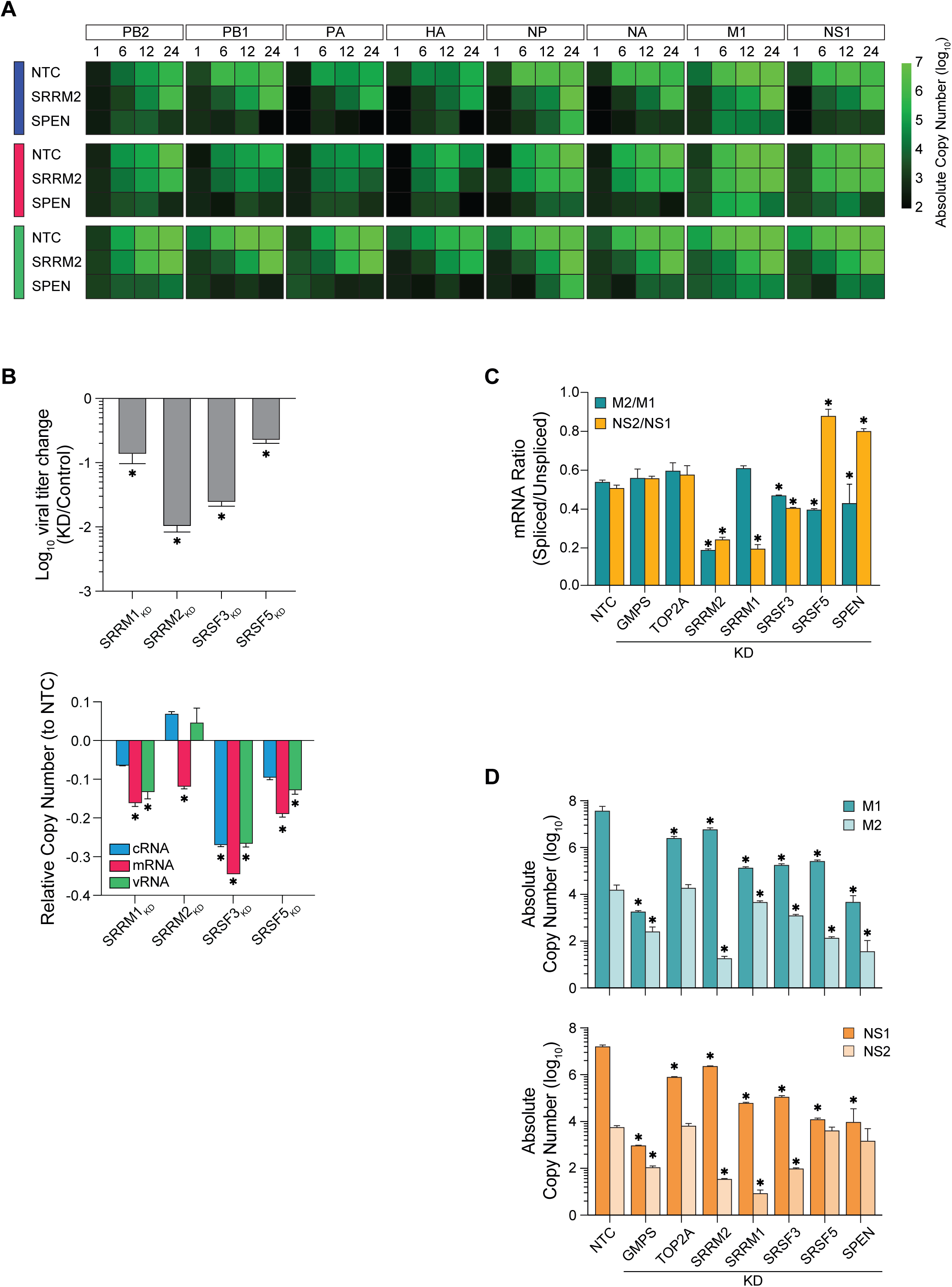
Dynamic regulation of IAV splicing by SR-proteins and SPEN. (A) Heatmap of IAV replication kinetics under wild-type, SRRM2, and SPEN knockdown conditions were monitored for each segment’s various RNA strands over a 24 h period (1, 6, 12, and 24 h). RNA absolute copy number is represented from low (black) to high (bright green). (B) Gene knockdown of SR-proteins, SRRM1, SRRM2, SRSF3, and SRSF5, negatively impact viral titer as measured by TCID_50_ assay (top) and viral RNA replication as measured for segment 1 (PB2) c-, m, and vRNA (bottom). (C) Ratio of unspliced to spliced viral mRNA for segments 7 and 8 (NS and M, respectively) (top). Ratio is calculated by dividing the spliced NS2/NEP or M2 by the spliced NS1 or M1 mRNA copy number, respectively. Absolute copy number of each mRNA splice variant (M1, M2, NS1, and NS2/NEP) within each knockdown condition. (D) Absolute copy number of IAV segments 7 and 8 unspliced (NS1 and M1, resplectively) and spliced (NS2/NEP and M2, respectively) within gene knockdown conditions. Error bars represent mean +/- SD, n = 3. * = p <= 0.05 (in B and C)

SRRM2 recognizes splicing elements on precursor RNA and organizes dynamic splicing condensates^96–98^. IAV has been shown to specifically utilize host nuclear speckles to promote the post-transcriptional splicing of its M1 mRNA^20^. Additionally, SRRM2 has been implicated in the co-localization and recruitment of SRSF proteins, often with SRRM2 within the core and SRSF2 at the extremities of speckle liquid condensates. Therefore, SRRM2, as a core scaffold protein of these speckles and a key splicing factor^96–98^, is highly likely to be directly co-opted by IAV to facilitate efficient viral mRNA processing as shown by our VIR-CLASP data.

As such, we wanted to investigate what the role of SRRM2, along with other SR-proteins within our dataset (SRRM1, SRSF3, SRSF5), and SPEN play in IAV splicing. Using primers targeting the unspliced M1 and NS1 and spliced, M2 and NS2/Nuclear Export Protein (NEP) forms of segments 7 and 8, respectively, we quantified the amount of each mRNA under knockdown conditions of each host factor. We show that knockdown of SRRM2, SRRM1, SRSF3, and SRSF5 reduced the viral titer by 100-, 10-, ∼90-, and ∼5-fold, respectively (Figure 6B, top). The effects on viral RNA species upon each knockdown varied. SRRM1, SRSF3, and SRSF5 reduced all three viral RNA species while SRRM2 reduced only viral mRNA (Figure 6B, bottom). This suggests that the spliceosomal complex and/or nuclear speckle condensates have some redundancy which allows for varied degrees of tolerance against loss/reduction in any one of these host proteins. IAV M1 and NS1 mRNA are spliced into various forms, including, M2, M3, M42, and NS2/NEP, respectively^17–20,137^. We demonstrate that knockdown of host factors unrelated to splicing machinery, such as GMPS and TOP2A, had no effect on the ratio of unspliced to spliced M and NS mRNA (Figure 6C). In contrast, SRRM2 knockdown resulted in significant decrease in spliced forms of both M and NS mRNA while, interestingly, SRRM1 reduced only NS2/NEP mRNA. SRSF3 knockdown decreased M2 and NS2/NEP, while SRSF5 depletion increased NS2/NEP and decreased M2 mRNA. This is supported by previous observations on SRSF5-related M1/M2 splicing^138^ and demonstrates that the mechanisms underlying IAV mRNA splicing are complex and dependent on the presence of host factors engaging with the viral mRNA. We further show that SPEN knockdown altered the ratios of spliced and unspliced M and NS mRNA (Figure 6C). The amount of spliced M2 mRNA significantly decreased while the spliced NS2/NEP mRNA significantly increased. This observation is interesting since NS1 is involved in autoregulation of its own mRNA splicing^139^. It has been reported that NS1, through engagement of m^6^A machinery, facilitates m^6^A modification of its own mRNA. This, in turn, prevents splicing of NS1 mRNA through competitive blockage of the SRSF3 binding site within NS1 mRNA by host YTHDC1 (enriched within our IAV interactome). Our opposite SPEN-related observation hints at the potential of SPEN influencing the proportion of IAV mRNA splice variants through its interactions with SR-proteins^136^ and m^6^A modification machinery^99^. This poses an interesting role for SPEN to add to its already known transcriptional regulatory function.

Our results indicate that GMPS, TOP2A, SRRM2, and SPEN are robust pro-viral host factors critical for IAV RNA genome replication. TOP2A appears to regulate viral replication processes involved in the synthesis of viral mRNA while GMPS and SPEN regulate processes involved in the production of viral c-, m-, and vRNA. Additionally, SPEN and SRRM2 (along with other SR-proteins) influence IAV splicing. However, it is difficult to delineate the effect on vRNA from that of c- and mRNA since vRNA synthesis is dependent on cRNA template which itself requires transcription by trans-vRdRp^50^. Therefore, any significant impact on viral m- or cRNA would likely translate into an impact on both c- and vRNA.

## DISCUSSION

Viruses are traditionally known to be sub-organismal parasites that hijack host cellular machinery for its own replication, but the precise molecular mechanisms remain under investigation. Leveraging advanced “-omics” technologies and the specific VIR-CLASP methodology, which uses 4SU-labeling to trace incoming genomic vRNA, we investigated the earliest host-virus interactions of IAV in human A549 lung epithelial cells. Within 1 hpi, the IAV vRNA is deployed and engages with over 600 host proteins, establishing a much broader interactome than previously understood, enriched in factors governing RNA metabolism, gene expression, and immunity. vRdRp components were not detected, likely since the viral polymerase is known to engage with host mRNA during the cap-snatching process^15,16,140–142^ which may have affected the crosslinking of the genome to the polymerase subunits. Additionally, the interaction between vRNA and the vRdRp complex is directed mainly through interaction of PA to the 5’-end of the vRNA while PB1 binding to PA is required for the PB1-vRNA interaction at this same locus^142,143^. This region of the vRNA is Uridine (U)-poor and Adenosine-rich, further explaining the lack of the vRdRp complex within the proteomics data due to the U-dependent nature of VIR-CLASP cross-linking^53,54,144,145^. The incoming IAV vRNA engages with upward of 600 host proteins within the first hour of entering a host cell of which ∼18% had been previously identified in RBP screens by eRIC^108^ and ivRIC^146^. Thus, the host-virus RNA interactome spans across a much larger group of proteins than previously described.

Assessment of the interactome revealed an enrichment of cellular processes involving RNA metabolism, gene expression and regulation, and immunity. Following initial screening by siRNA knockdown of host factors representative of each process, we selected GMPS, TOP2A, SRRM2, and SPEN for further validation. We demonstrated upwards of a 100- to 10,000-fold decrease in viral titer and decreases in viral RNA species (v-, c, and mRNA) for all gene segments as early as 6 hpi, which varied for each target. The effects observed imply a different role for each host protein during the viral replication cycle and the importance of RBP interactions with the incoming vRNA prior to cRNA and mRNA production.

GMPS is an interesting candidate for IAV therapeutics since both gene knockdown and pharmacological inhibition of GMPS resulted in negative impacts on IAV replication. GMPS catalyzes the conversion of xanthosine monophosphate to GMP using ATP and l-glutamine^101,102^. This reaction is vital for providing GMP essential for DNA/RNA replication and transcription. Beyond its primary metabolic role, GMPS exhibits "moonlighting" functions that directly influence gene regulation through mediating p53 stabilization^147,148^, hyperactivation of ubiquitin-specific protease 7 (USP7)^148^ - deubiquitylating histone H2B^149^. This interaction regulates Epstein-Barr Virus replication via the viral EBNA1-USP7-GMPS interaction^150^. GMPS has recently been demonstrated to also have RNA-binding activity via various methodologies^103–108^. Thus, in the context of IAV infection, purine biosynthesis is of particular interest since the vRdRp initiates replication through the characteristic incorporation of a pppApG dinucleotide – essential for replication to start^111,112^. Importantly, this activity is not derived from a host protein. Alteration of cellular GMP levels could have wide-spread effects on several cellular processes, including RNA synthesis of host transcripts. However, our results show a more specific viral effect upon GMPS depletion or catalytic inhibition, at least within the time course of the assay – and suggesting that GMPS activity may be acutely required for the *local synthesis* of GMP at viral replication sites. This is suggested by the inability of GMP supplementation to rescue IAV replication and initiation under GMPS depleted conditions despite the salvage pathway being intact.

Cell culture medium does not provide a source of purine bases, but FBS does^151^, supplying the salvage pathway with the necessary components for GMP synthesis. This pathway may utilize either 1) convert guanine to GMP directly or 2) first convert guanine to guanosine prior to conversion to GMP^109^. Therefore, when the *de novo* pathway is impaired, the salvage pathway may compensate in several ways^152^. This pathway is exploited by the pharmacological agent Ribavirin, which impede IAV replication through inhibition of IMP dehydrogenase^153^, an enzyme upstream of GMPS, within the *de novo* biosynthetic pathway of GTP. Although, having a pool of GMP available may not be sufficient to recover viral replication, for example, HIV infection is delayed when ablating the *de novo* biosynthesis pathway using ribonucleotide reductase inhibitors^154^. Ribavirin has been demonstrated to reduce vRNP synthesis by 50% at 25 µM and 95% at 100 µM ribavirin in cell culture. However, IAV has shown resistance to ribavirin impacting its feasibility as clinical intervention^155^. Our study identified target proteins with a comparable impact on IAV titer to that of a known anti-virals. Dysregulation of GMPS is also associated with acute myeloid leukemias and cervical cancer, further highlighting the expanding role of GMPS in host cellular regulation and health – providing a potential candidate for novel therapeutics as not only an anti-viral, but also in treatments for cancer.

TOP2A, along with other topoisomerases, significantly impacts transcription^92^ through its close relationship with RNA Pol II by promoting productive transcription elongation through facilitating the release of paused RNA Pol II^91^. TOP2A, and TOP1^156^, interactions have been expanded to RNA binding^65,67,103,105–107,113–116^ and functions involving RNA-dependent proteins, which may modulate its DNA relaxation activity. Given IAV’s nuclear replication cycle and its reliance on host RNA Pol II for viral transcription^15,93–95^, TOP2A’s function in maintaining a permissive chromatin environment, and its involvement in promoter-proximal pausing of RNA Pol II in certain contexts, may be co-opted by the virus^117^. TOP2A knockdown showed a negative effect on viral c- and mRNA (although the cRNA levels recover over time) (Figure 5A and see also Figure S5A). Our data further demonstrates that therapeutic inhibition of TOP2A altered the dynamics of viral RNA species. This is likely due to the mechanism of inhibition which creates a rigor complex between TOP2A and its substrate. This complex would prevent the replication of all viral RNA species through physical obstruction of the vRdRp and further points to TOP2A directly interacting with the incoming vRNA. Furthermore, knockdown and pharmacological inhibition of TOP2A negatively influenced the degree of 5’-capped viral mRNA relative to control. The under-capping of viral mRNA would interfere with mRNA nuclear export and result in destabilization of the RNA. It is possible that TOP2A engages with vRNA for the purpose of assisting in cap-snatching through its interweavement with host RNA Pol II. TOP2A closely associates with the host DNA during transcription events and has been shown to instigate RNA Pol II stalling. IAV RdRp is known to interact with the CTD of RNA Pol II and is thought to use this interaction to bring the vRNA template near the nascent strand of host pre-mRNA. Thus, the earlier reduction in cRNA levels for segments 3 through 6 may be due to the unique situation where IAV mRNA (following cap-snatching), is subject to poly-A signal read-through – generating hybrid capped cRNA^157^ which may be reduced when TOP2A is depleted. Together TOP2A and vRdRp may thus facilitate the cap-snatching process of IAV.

SRRM2 contains an N-terminal RRM for direct RNA binding, two large serine/arginine-rich (RS) domains that mediate protein-RNA interactions, and IDRs that facilitate protein-protein interactions^96^. SRRM2 is known to be involved in nuclear speck formation and acts as core component of the spliceosome through interaction with other splicing factors such as SON and SRSF2^96–98,158^. Additionally, SRRM2 forms splicing co-activator complexes with its smaller, IDR-rich homologue, SRRM1^96^. SON and SRRM2 form separate nuclear speckle sub-compartments, each containing a different subset of proteins. SRRM2 also oligomerizes through its RS domains driving phase separation, facilitated by RNA interaction^98^. Nuclear speckles have also been demonstrated to recruit host machinery involved in various gene expression and regulation pathways such as transcription (LARP7), elongation (CDK12), and m^6^A RNA modification (WTAP, METTL3, RBM15, YTHDC1)^159–161^ (many of which were present in our IAV interactome). Thus, nuclear speckles are not only driving splicing but also recruit machinery to the RNA site under conduction by SR-proteins. Since IAV is known to utilize host splicing machinery to splice some of its viral mRNA (M and NS)^20,45,50^, we sought to investigate the effects our candidate host factors such as SRRM2 have on IAV splicing.

SPEN contains N-terminal RRMs and a C-terminal SPOC domain which has been documented to recognize phosphorylated serine motifs in m^6^A reader and writer proteins such as FMR1^162^. SPEN plays a role in recruitment of m^6^A machinery, as demonstrated in the context of transcriptional regulation of *Xist* through SPEN-mediated RBM15 and ZC3H13 recruitment^99^. Our IAV VIR-CLASP interactome captured several host m^6^Amachinery components such as FMR1^162,163^, RBM15, RBM15B, and ZC3H13^164^. IAV RNA is known to be post-transcriptionally modified, including m^6^A modifications^165,166^. Studies report that IAV m^6^A modification occur on both the minus- and plus-sense RNA^167^ and modification of vRNA promotes engagement with the vRdRp^168^. Deficient vRNA m^6^A modification impairs overall replication of IAV^167^. Our data show that SPEN knockdown impaired replication of all viral segments and RNA species (with some skew toward mRNA). Studies have shown that IAV NS1 regulates its own splicing through interaction with METTL3/14 and exploitation of m^6^A modification on NS1 mRNA^138,139,166^. Although the m^6^A modification role of SPEN is an intriguing model warranting further investigation, for the scope of this study we focused on the potential role of SPEN in splicing. SPEN, through its interactions with RNA and m^6^A host machinery, may therefore influence the deposition of m^6^A to IAV RNA which in turn influences the RNA fate. SPEN involvement in m^6^A modification of RNA is further supported by studies demonstrating WTAP (an interactor of SPEN) recruitment of the m^6^A “writer” complex to splice sites via interactions with the METTL3/METTL14 heterodimer^169,170^. RBM15/15B have also been implicated in METTL3/METTL14 recruitment to RNA^171^. In METTL3 expression defective cells, IAV replication was found to decrease approximately 8-fold^167^ – although significant, implies some level of redundancy may be involved in the absence of METTL3.

Recent research has shown that the Repeat A domain in *Xist*, although critical for SPEN recruitment, does not directly interact with SPEN but rather SR-proteins^136^. SR-proteins, such as SRSF1, directly interacts with SPEN and aid in its recruitment to *Xist* Repeat A domain. This is an intriguing development which may imply that SR-proteins, like SRSF3 found in our IAV interactome, not only direct splicing events, but also recruit SPEN and its transcriptional regulatory machinery to the nuclear speckles formed under the direction of SRRM1, SRRM2, and SON. We demonstrated that SR-proteins dynamically influence the splicing of IAV segments 7 and 8. Altered splicing events lead to changes in the ratio of viral proteins and affect viral protein functions such as NS2/NEP which is responsible for nuclear export of viral mRNA^172^ – especially the intron-containing HA, M, and NS mRNA. With impaired splicing, IAV replication, nuclear egress, and eventual virion assembly and release from the cell will be defective. Thus, the role and the molecular makeup of nuclear speckles need further investigation to elucidate whether the condensates formed by SR-proteins during splicing events are also utilized by viruses, like IAV, to assemble into viral replisomes containing all the necessary host machinery to drive rapid and successful viral replication. Together, the SR-proteins and SPEN play important roles in splicing of IAV, which may be directed through the assembly of the spliceosomal complex and/or m^6^A modification of viral transcripts.

Our research demonstrates how the incoming IAV genome is more than merely a template for viral cRNA and mRNA transcription; it also serves as a central functional scaffold for recruiting and coordinating a vast network of critical host factors. This recruitment strategy ensures the timely and efficient assembly of the molecular machinery required for subsequent viral replication. By engaging a suite of essential host proteins – including splicing factors (like SR-proteins), transcriptional regulators, and nucleotide biosynthetic enzymes – the vRNA successfully orchestrates the early cellular environment to favor viral replication. This nuanced role of post-entry RBPs in regulating IAV through direct interaction with the incoming genome highlights their potential as novel antiviral targets. Moreover, the apparent conservation of such interactions across viral strains and species underscores the importance of exploring these early viral-host interactions for broad antiviral strategies.

## RESOURCE AVAILABILITY

### Lead contact

Manuel Ascano Jr.

### Materials availability

Further information and requests for resources and reagents should be directed to and will be fulfilled by the Lead Contact, Manuel Ascano.

### Data and code availability

The mass spectrometry proteomics data have been deposited to the ProteomeXchange Consortium (<u>Deutsch et al., 2017</u>) via the PRIDE (<u>Perez-Riverol et al., 2019</u>) partner repository with the dataset identifier PRIDE: *in progress*. All code used for proteomic analysis and figure generation is accessible at https://github.com/Ascano-Lab.

## ACKNOWLEDGMENTS

We would like to thank Dr. C. P. Plamen and Vanderbilt Institute of Chemical Biology Chemical Synthesis Core for 4SU synthesis; the Mass Spectrometry Research Center Proteomics Core Laboratory (Vanderbilt University), in particular Dr. Kristie Lindsey Rose and Dr. Michael Leser for mass spectrometry analysis; Dr. Suman Das for guidance on IAV culture methodology; Dr. Lars Plate for discussion in the MS analysis. Finally, we would like to thank members of the Ascano laboratory for their support, collegiality, and critical review of the manuscript. This work was supported by the NIH (5R35GM119569) and NIH T32 training grant for their support (training grant 5T32AI138932-04 and 5T32AI138932-05 to S.C.).

## AUTHOR CONTRIBUTIONS

Conceptualization, S.C. and M.A.; methodology, S.C. and M.A.; Investigation, S.C., D.P., C.L.W., J.L.P.; software and data curation, S.C., A.A. and S.L.L., writing—original draft, S.C; writing—review & editing, S.C and M.A.; funding acquisition, M.A.; resources, K.L.R, M.A.; supervision, M.A.

## DECLARATION OF GENERATIVE AI AND AI-ASSISTED TECHNOLOGIES

The authors did not use AI technologies.

## SUPPLEMENTAL INFORMATION

**Document S1. Figures S1–S4.**

**Table S1. IAV VIR-CLASP Proteomics Analysis, Related to Figure 1**.

**Table S2. Databases used for computational comparisons, Related to Figure 2**.

**Table S3. List of RBPs in IAV interactome, Related to Figure 2**.

**Table S4. Functional analysis of IAV interactome, Related to Figure 2**.

**Table S5. Final list of IAV interactome hits and pathway analysis, Related to Figure 3A**.

**Table S6. Primers and siRNAs used in this study, related to the STAR Methods.**

## STAR★METHODS

### KEY RESOURCES TABLE

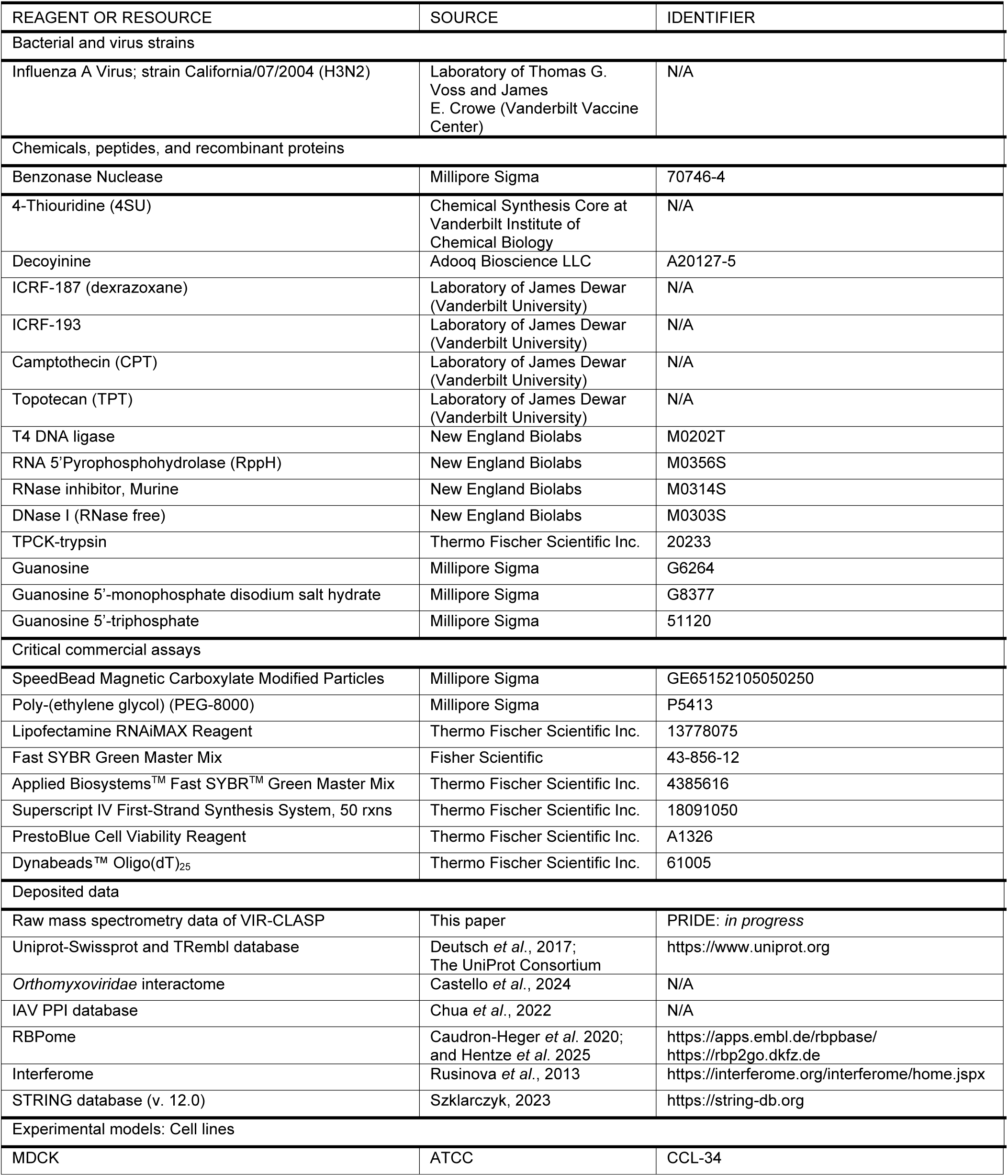
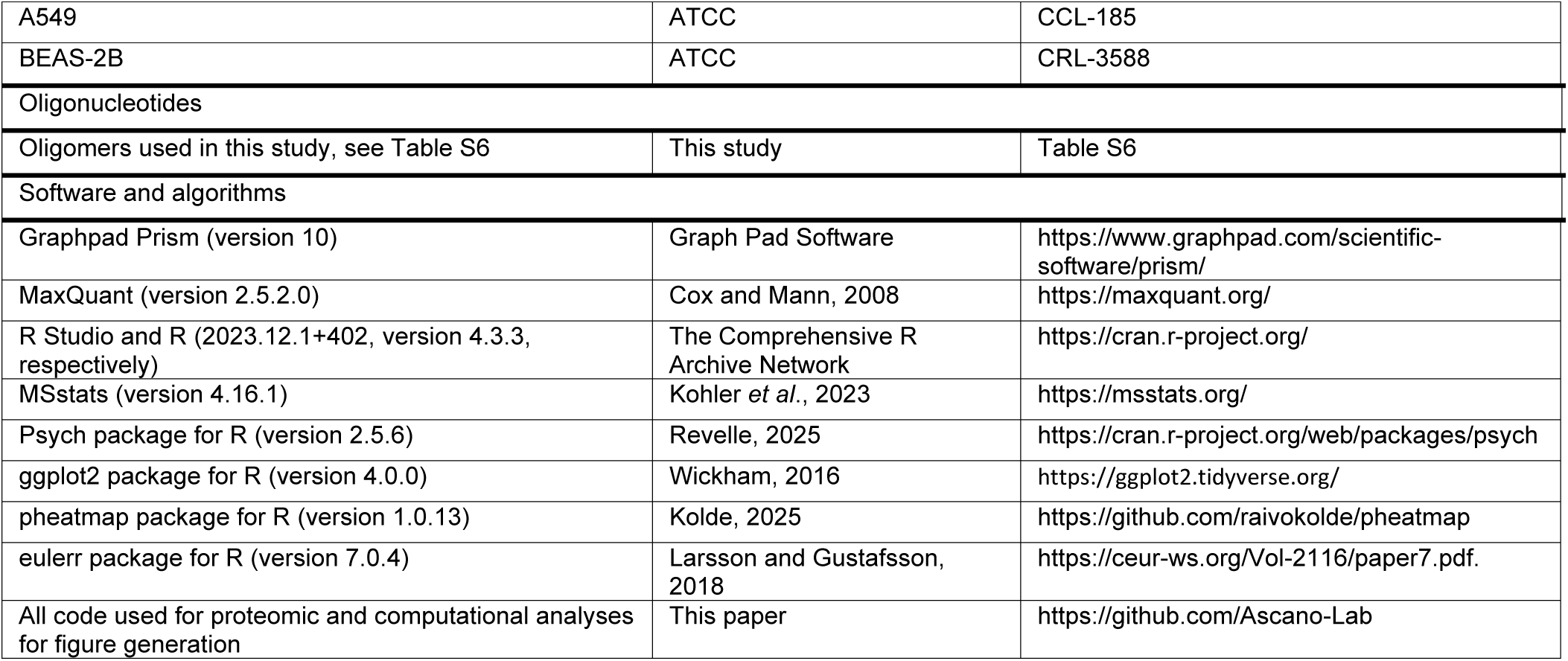

### EXPERIMENTAL MODEL

#### Cell lines and culture

All cell lines were obtained from ATCC. A549 cells (male) were maintained in DMEM (Gibco) supplemented to contain 10% fetal bovine serum (FBS from Peak Serum) and 2 mM L-Glutamine (Gibco). MDCK (female canine) were maintained in MEM without glucose supplemented with 7.5% FBS and GlutaMAX (Gibco). All cultures were maintained in a 5% CO_2_, humidified incubator at 37 °C.

#### Viral Culture

Influenza A virus (IAV; strain: California/7/2004 (H3N2) was provided by Thomas G. Voss and James E. Crowe (Vanderbilt Vaccine Center). For 4SU incorporation into the viral genome 0.1 mM 4SU was added to MDCK culture medium on the day of subculture, one day prior to infection with IAV. On the day of infection, the medium was exchanged for IAV infection medium (MEM supplemented with GlutaMAX and 1 µg/mL TPCK-trypsin) containing MOI 0.1 of IAV. The virus was allowed to infect the cells for 1 h followed by removal of the inoculum and washing the monolayer with MEM medium. Fresh infection medium was added and supplemented with 1 mM 4SU.

Virus stocks were purified by ultracentrifugation of clarified supernatants through a 30% sucrose cushion in NTC buffer (50 mM Tris-HCl [pH 7.2], 0.1 M NaCl, and 0.1 M CaCl_2_) at ∼121,000 × g for 3 h in a Beckman SW32Ti rotor and then re-suspended in virus infection medium, aliquoted, and stored at −80 °C. A549 and BEAS-2B infections used VIR-CLASP infection medium (DMEM supplemented with 0.3% BSA, 10 mM HEPES, and 0.5 µg/mL TPCK-trypsin). Viral titer was determined by TCID_50_ assay in MDCK cells.

### METHOD DETAILS

#### Viral Titer Determination (TCID_50_)

Cell culture medium was collected from infected cells at the desired time post-infection. One day prior to collection, MDCK cells were seeded into 96-well plates at 2 million cells total per plate. On the day of infection medium collection, the MDCK cells were washed three times with MEM (no supplements) and serial dilutions (10-fold) of the infected culture medium prepared in viral infection medium. Dilutions were added to wells containing washed MDCK cells and incubated under culture condition as described above for 100 h. After incubation, MDCK cells were washed with PBS and 0.1% crystal violet stain containing fixative added to each well and incubated at room temperature for 30 minutes prior to washing in PBS and assessing of virus-positive wells. Viral titer was calculated using the Reed-Muench method^173^.

### VIR-CLASP

#### Viral infection and UV crosslinking

Cells were infected with 4SU-labelled virus (MOI 1000) by adsorption for 1 h at 4°C followed by washing uninfected virus away with cold VIR-CLASP infection medium. The infected cells were incubated for at 37°C for 1 h, washed with cold PBS, and irradiated on ice with 365 nm ultraviolet light (0.6 J/cm^2^ x 2 times) in a Stratalinker 2400 (Stratagene). Cells were scraped off in 2.5 ml PBS per plate, transferred to a centrifuge tube, the plate washed with an additional 2.5 mL cold PBS to collect remaining cells. The cell suspension was then centrifuged at 300 x *g* for 5 minutes, PBS removed, and cell pellets used for CLASP procedure as described previously^145^.

#### CLASP

Cells were lysed in denaturation buffer (50 mM Tris–HCl, pH 6.8, 10% glycerol, 2.5% SDS, 0.66% NP-40), incubated for 10 min at 95 °C and subsequently slowly cooled to 25 °C. Crosslinked RNA-protein complexes were purified by Solid-Phase Reversible Immobilization (SPRI) (Hawkins et al., 1994) beads (GE Healthcare) under denaturing SPRI buffer (30 mM Tris–HCl, pH 6.8, 6% glycerol, 1.5% SDS, 0.4% NP-40, 1 M NaCl, 8% PEG-8000). To each sample, 0.66x (e.g. 660 μL of beads for 1 mL of sample) of SPRI beads (1 mg/mL SPRI beads in 10 mM Tris-HCl, pH 8.0, 1 M NaCl, 18% PEG-8000, 1 mM EDTA and 0.055% Tween 20) were added, and samples were incubated at room temperature for 10 minutes. The SPRI beads and complexes were washed 2 times with denaturing SPRI buffer. The crosslinked RNA-protein complexes were eluted for 5 min at 37 °C in denaturation buffer (lysis buffer). To reduce non-specific binding on the beads, SPRI purification was repeated. To digest RNA from crosslinked RNA-protein complexes, an equal volume of 4x Benzonase buffer (80 mM Tris-HCl, pH 7.5, 600 mM NaCl, 20 mM MgCl2, 4 mM DTT, 40% Glycerol) and 2x volume of water were added to eluted samples, followed by the addition of Benzonase (EMD Millipore) to a final concentration of 50 U/mL, and incubation for 2 h at 37°C. The final eluted and Benzonase-treated proteins were precipitated by methanol and chloroform and then re-suspended in 2x NuPAGE LDS Sample Buffer (Thermo Fisher Scientific, cat# NP0007) with 50 mM DTT for SDS-PAGE and silver staining or 100 mM TEAA containing 1% SDS for mass spectrometry analysis.

#### Mass Spectrometric Analysis

Protein samples were brought to 5% SDS, prepared by S-Trap (ProtiFi) digestion, and analyzed by LC-coupled tandem mass spectrometry (LC-MS/MS) as previously described^174^. Proteins were initially reduced with 10 mM TCEP, alkylated with 20 mM iodoacetamide, and aqueous phosphoric acid was added at a final concentration of 2.5% followed by S-trap binding buffer (90% methanol in 100 mM TEAB) at 6 times the sample volume. Samples were loaded on S-Trap micro columns, washed with S-trap binding buffer, and were digested with 1 ug trypsin gold (Promega) in 50 mM TEAB, pH 8.0, for 1 h at 47 °C. Peptides were eluted by serial addition of 40 µL each of 50 mM TEAB, 0.2% formic acid, and 35 µL of 0.2% formic acid in 50% acetonitrile. Eluted peptides were dried and resuspended in aqueous 0.2% formic acid. For LC-MS/MS, peptides were loaded onto a C18 reverse phase analytical column using a Dionex Ultimate 3000 nanoLC and autosampler and were gradient eluted using a 90 min gradient. The gradient consisted of the following: 1-77 min, 2–38% B; 77-82 min, 38–95% B; 82–83 min, 95% B; 83-84 min, 95–2% B; 84-90 min (column re-equilibration), 2% B. Peptides were analyzed using a data-dependent method on an Orbitrap Exploris 240 mass spectrometer (Thermo Scientific), equipped with a nanoelectrospray ionization source. The instrument method consisted of MS1 using an MS AGC target value of 3 × 10^6^, followed by 20 MS/MS scans of the most abundant ions detected in the preceding MS scan. The intensity threshold for triggering data-dependent scans was set to 1 × 10^4^, the MS2 AGC target was set to 1 × 10^5^, dynamic exclusion was enabled (15 s), and HCD collision energy was 30 NCE.

#### Bioinformatics analysis of Mass Spectrometry data

Raw data files from the LC-MS/MS instrument were processed and searched using MaxQuant (v2.5.2.0)^175,176^ to generate peak lists and identify peptide-spectrum matches. Searches were performed using a Uniprot/Swissprot database for *Homo sapiens* (proteome ID: UP000005640) and *Canine lupus familiaris* (proteome ID: UP000805418) including both reviewed (Swissprot) and unreviewed (TrEMBL) proteins (downloaded on 22 April 2024), with added sequences for Benzonase nuclease (Uniprot Accession: P13717), and for IAV (H3N2) proteins (PB2, PB1, PA, HA, NP, NA, M1, M2, NS1, NS2/NEP) from strain A/CA/07/04. The search parameters for Andromeda included full tryptic specificity, two missed cleavages allowed, carbamidomethyl (C) fixed modification, and acetylation (N terminal) variable modification. Match between runs was selected, and LFQ normalization with iBAQ identification algorithm.

All other settings used were default, resulting in a protein FDR of < 0.01 for each dataset. To define the set of pre-replicated IAV interacting proteins, we used the MSstats v 4.16.1 R package^56^. Potential contaminants were kept, proteins identified by only one peptide removed, and protein identification performed using unique peptides. Quantile normalization was applied to remove systematic bias between MS runs, model-based imputation performed for censored missing values using accelerated failure time model, and remaining parameters were set to default. For statistical comparison, VIR-CLASP +4SU samples were compared to the corresponding input +4SU sample. Adjusted P values were computed using Student’s t-test with Benjamini-Hochberg correction. Proteins with an adjusted p-value ≤ 0.05 and a log_2_ fold-change > one were classified as enriched VIR-CLASP proteins (IAV RNA Interactome). Correlations between replicates were performed using Pearson correlation.

#### Functional enrichment analyses

Functional enrichment clusters of Gene Ontology terms in protein-protein association network were retrieved and clustered STRING v12.0 database. Enriched Reactome pathways and InterPro classifications were queried using STRING v12.0 database and applying default settings. Statistical background for enrichment analysis was performed using the whole genome.

#### siRNA knockdown

A549 cells were seeded in 24-well plates at 1.5 × 10^5^ cells/well and incubated for 24 h prior to transfection. siRNAs (listed in Table S6) were transfected at a final concentration of 20 nM using Lipofectamine® RNAiMAX (Invitrogen) according to the manufacturer’s instructions. Cells were incubated at 37 °C and 5% CO_2_ for 72 h before infection with IAV at an MOI of 0.1.

#### Inhibition assays

GMPS inhibitor, decoyinine (Adooq Biosciences), was reconstituted in 100% DMSO as 17.9 M stocks. Cells were treated with decoyinine (1.79 µM), ICRF-187 (10 µM), ICRF-193 (0.5 µM), CPT (0.68 µM), TPT (1 µM), diluted in infection medium, either 24 or 72 h prior to infection with IAV at MOI 0.1 or cell viability assessment as described below. Fresh inhibitors were supplemented during IAV infection before culture medium and cell lysate collection at 24 hpi. IAV titer and RNA were assessed as described above.

#### Cell viability assay

A549 cells were seeded in 96-well plates at 1.5 x 10^5^ cell/well and treated with the above listed inhibitor concentrations. Stock solutions were diluted in infection media prior to cell treatment. DMSO was used as a vehicle control. After 72 h of treatment, cell viability was assessed using PrestoBlue reagent (Thermo Scientific) according to the manufacturer’s instructions.

#### RNA extraction and RT–qPCR analysis

RNA was collected from infected cells using Trizol (Ambion). The RNA concentration was determined using a NanoDrop 2000 (ThermoFisher). Equal amounts of total RNA (1 µg) were reverse-transcribed using SuperScript IV (ThermoFisher) with random hexamer primers (for total RNA) or IAV strand-specific primers. Quantitative PCR reactions were performed using FastSYBR Green Plus Master Mix (Applied Biosystems) using a QuantStudio™ 3 Real-Time PCR instrument (Applied Biosystems). Oligonucleotides used in this study are listed in Table S6. Target Ct values were normalized to *TUBA1A* Ct values and used to calculate ΔCt. Relative mRNA expression of target genes was then calculated using the ΔΔCt method (2^ΔΔCt^). Absolute quantification of IAV RNA was performed using a standard curve of a plasmid containing a cDNA copy of region of the IAV strain A/CA/07/04 (H3N2) (pUCIDT-Amp, IDT) was used. Viral copy number was calculated based on absolute quantification using the standard curve and relative viral copy number was based on control (NTC) values.

For segment-specific assays a time-course of 1, 6, 12, and 24 hpi was used with RNA collected at each time as described above, prior to RT-qPCR. For the RT reactions, strand-specific primers were used targeting the universally conserved 5’-Uni-12 sequence present in all eight IAV segments (5’-AGCG(A/G)AAGCAGG-3’,), the 3’-Uni-13 sequence (5.-AGTAGAAACAAGGTCG-3’), or sequences targeting the poly-A tail (Table S6).

For splicing assays reverse transcription was performed as above using oligo-dT (Invitrogen) primers. Primers targeting specific regions for M1 and NS1 or M2 and NS2/NEP were used to distinguish between unspliced and spliced versions of segment 7 and 8, respectively (Table S6).

#### Cap-interference RT-qPCR

The proportion of viral mRNA containing 5’-cap moieties were assessed by cap-interference RT-qPCR. First, total mRNA was extracted from infected A549 cells at the stated MOI and time-point using oligo-d(T)_25_ (Invitrogen) beads as per the manufacturer’s instructions. Next, the mRNA was treated with or without RppH (25 U, NEB) digestion of 5’-cap as per the manufacturer’s instructions for decapping eukaryotic mRNA. 100 µM DNA splint and RNA adapter were mixed in a ratio of 1:2 (splint:adapter) and incubated at 60 °C for 5 min. Adapter-splint probes were added to mRNA and annealed in a step-wise temperature sequence: 70 °C, 5 min; 60 °C, 5 min; 42 °C, 5 min; 25 °C, 5 min prior to addition of T4 DNA ligase (400 U, NEB) and incubation at 15 °C for 18 h. DNA splint was removed by digestion with DNase I (10 U, NEB) and inactivated. Finally, cDNA was generated using primers targeting the adapter sequence (Table S6) as above. To calculate the proportion of capped mRNA, the RppH-treated or control Ct values were corrected to account for differences in total viral RNA load using a total viral RNA control (corrected Ct). The corrected Ct values were normalized using *TUBA1A* Ct values (delta corrected Ct). The proportion of capped transcripts are finally calculated by comparing the delta corrected Ct of RppH-treated to control (delta-delta corrected Ct).

#### Reverse-Phase HPLC

For the estimation of free intracellular nucleotides within A549 cells under described conditions, 1 mL of TriZol reagent was added to washed cells prior to phase separation using chloroform and collection of the aqueous phase. The RNA and free nucleotide/nucleoside-containing aqueous solution was passed through a miRNeasy (Qiagen) column to remove total RNA and the *flow-through* collected. Next, the samples were evaporated under vacuum until the volume reached ∼50 µL. The concentrated samples were then added to nucleotide-precipitation solution (PS) (100 mM sodium acetate, 100 mM lanthanum chloride, and 0.6 mg/mL sodium phosphate mono-basic, pH 4.0) and nucleotide precipitated as described previously^177^. Final pellets were resuspended in 55 µL 0.5 M HCl and analysed by RP-HPLC^178^. Briefly, 50 µL of samples were injected and separated using a C18 column by gradient elution using solvent A (20 mM ammonium acetate, pH 5.4) and solvent B (100% methanol). The gradient started at 100% solvent A, ramped up to 25% solvent B over 16 minutes, ramped back down to 0% solvent B in 1 minute for 13 minutes (30 min total method time) at a flow rate of 0.5 mL/min and detection at 260 nm.

### QUANTIFICATION AND STATISTICAL ANALYSIS

Using the GraphPad PRISM 10 software, a two-tailed Student’s t-test and one-way ANOVA (Tukey) were used for statistical analysis of all data presented except mass-spectrometry and bioinformatics analysis. Numbers of biological replicates of assays (n) are provided and defined within the corresponding figures or figure legends. Error bars shown in the Figures represent means ± SD.

