## Supplemental Figures for "The Incoming Influenza Genome Assembles a Host RBP Network that Orchestrates Viral RNA Synthesis"

**Supplemental Figure 1. IAV replication kinetics in A549 and BEAS-2B cells.**

(A) Schematic representation of viral entry differences (left) and IAV intracellular copy number of segment 1 following unequal (top, right) and equal (bottom right) entry.

(B) Quantification of IAV RNA for each segment within entry-normalized A549 and BEAS-2B cells over 72 h time-course (1, 6, 24, 48, and 72 hpi).

(C) Pearson correlation between VIR-CLASP samples. Numbers represent Pearson correlation coefficients; histograms represent distribution of LFQ intensities.

Error bars represent mean ± SD, n = 3 (for A and B).

**Supplemental Figure 2. RBP classifications within IAV interactome**

(A) The proteins within the RBPome and IAV interactome were assessed by Pfam identifiers to distinguish between conventional (cRBP), non-conventional (ncRBP), and presumed (pRBPs) RBPs.

**Supplemental Figure 3. Candidate protein network and knockdown**

(A) Protein-protein association network of filtered IAV interactome host proteins retrieved from STRING. Each circle (node) represents an individual protein with lines indicating interactions with another protein within the network. Colored nodes indicate proteins involved in RNA metabolism (blue), infectious disease and the immune system (greens), gene expression (red), or regulation of gene expression (orange).

(B) siRNA knockdown of IAV RNA interactome proteins in A549 cells followed by RT-qPCR to measure mRNA expression and evaluate knockdown efficiency. Data is normalized to *TUBA1A* using $\Delta\Delta C_t$ method and is relative to non-template control.

Error bars represent mean ± SD, n = 3 (for B).

**Supplemental Figure 4. Strand-specific replication under GMPS knockdown**

(A) Schematic representation of strand-specific RT-qPCR. The IAV genome shares universal sequences within all its segments (Uni-12 and Uni-13). Primers targeting these universal sequences generate cDNA of each strand (v- or cRNA). Viral mRNA contains a poly-A tail which is used to generate cDNA of viral mRNA. Each segment is then quantified using primers targeting each segment.

(B) Relative copy number of each strand for each IAV segment over 24 h time course (1, 6, 12, and 24 hpi). Control IAV-infected cells are in dotted lines with GMPS knockdown in solid.

(C) Cell viability upon decoyinine treatment.

Error bars represent mean ± SD, n = 3 (for B and C).

**Supplemental Figure 5. Strand-specific replication under TOP2A knockdown.**

(A) Relative copy number of each strand for each IAV segment over 24 h time course (1, 6, 12, and 24 hpi). Control IAV-infected cells are in dotted lines with TOP2A knockdown in solid.

(B) Cell viability upon topoisomerase inhibitor treatment was measured following 72 h by PrestoBlue quantification assay (see text).

(C) Schematic representation of cap-interference RT-qPCR method. Viral mRNA (collected by oligo-dT pull-down) is treated with RppH or not. The resultant RNA is ligated to an RNA adapter-DNA splint oligo, where the DNA splint has a sequence specific to viral mRNA. The DNA splint is digested away and the adapter sequence used for RT-PCR followed by IAV-specific qPCR. The ratio between RppH-treated and untreated RNA, normalized to total, is used to represent the proportion of capped transcripts.

Error bars represent mean ± SD, n = 3 (for A and B).

**Supplemental Figure 6. Strand-specific replication under SRRM2 or SPEN knockdown.**

(A) Relative copy number of each strand for each IAV segment over 24 h time course (1, 6, 12, and 24 hpi). Control IAV-infected cells are in dotted lines with SRRM2 knockdown in solid.

(B) Relative copy number of each strand for each IAV segment over 24 h time course (1, 6, 12, and 24 hpi). Control IAV-infected cells are in dotted lines with SPEN knockdown in solid.

Error bars represent mean ± SD, n = 3 (for A and B).

### Figure S1.

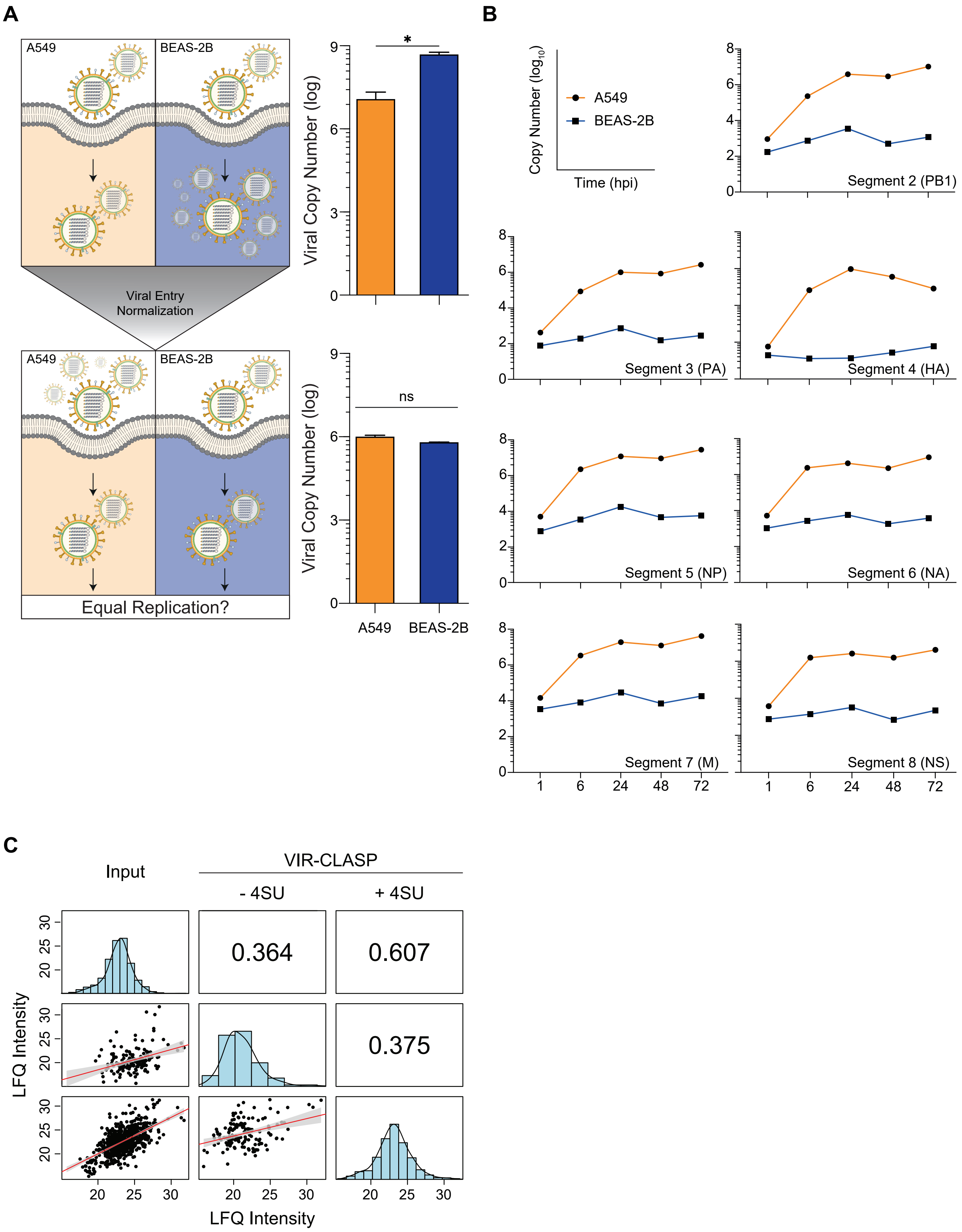
A
A549
BEAS-2B
Viral Entry
Normalization
A549
BEAS-2B
Equal Replication?
Viral Copy Number (log)
*
ns
A549
BEAS-2B
B
Copy Number (log10)
Time (hpi)
A549
BEAS-2B
Segment 2 (PB1)
Segment 3 (PA)
Segment 4 (HA)
Segment 5 (NP)
Segment 6 (NA)
Segment 7 (M)
Segment 8 (NS)
C
Input
VIR-CLASP
- 4SU
+ 4SU
0.364
0.607
0.375
LFQ Intensity
LFQ Intensity

### Figure S2.

**A**

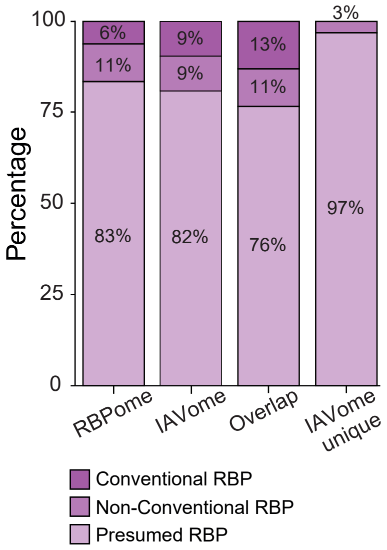

Percentage
100
75
50
25
0
6%
11%
83%
9%
9%
82%
13%
11%
76%
3%
97%
RBPome
IAVome
Overlap
IAVome unique
Conventional RBP
Non-Conventional RBP
Presumed RBP

Figure S3.

A

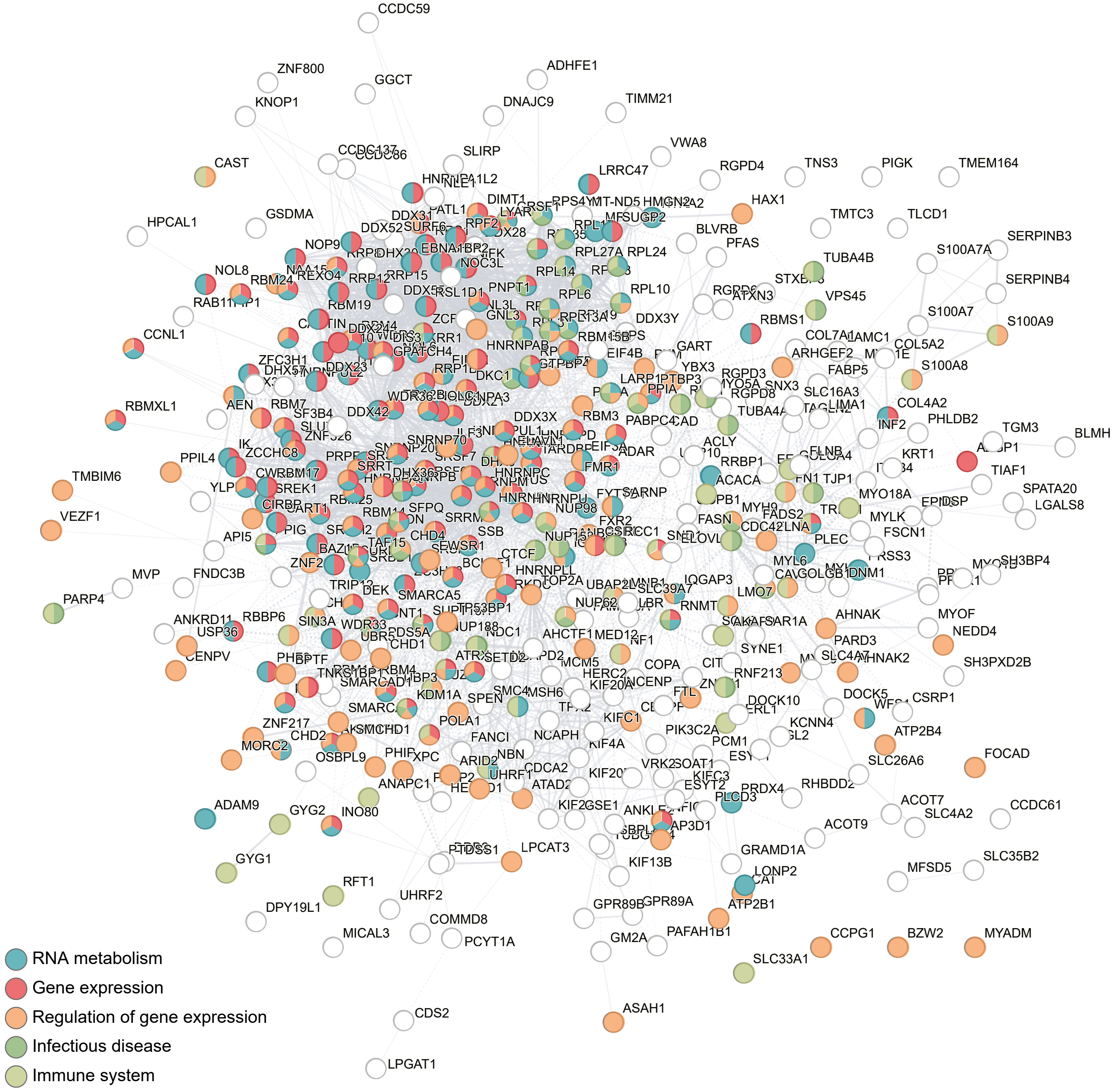
CCDC59
ZNF800
GGCT
ADHFE1
KNOP1
DNAJC9
TIMM21
VWA8
CAST
SLIRP
LRRC47
RGPD4
TNS3
PIGK
TMEM164
HPCAL1
GSDMA
HAX1
TMTC3
TLCD1
SERPINB3
S100A7A
TUBA4B
SERPINB4
VPS45
S100A7
S100A9
S100A8
CCNL1
RBMXL1
COL4A2
TGM3
BLMH
PPIL4
TMBIM6
VEZF1
TIAF1
SPATA20
LGALS8
MVP
FNDC3B
PARP4
MYOF
NEDD4
SH3PXD2B
CSRP1
ATP2B4
FOCAD
SLC26A6
RHBDD2
ACOT7
CCDC61
SLC4A2
ACOT9
GRAMD1A
SLC35B2
MFSD5
ADAM9
GYG2
INO80
GYG1
RFT1
DPY19L1
UHRF2
MICAL3
COMMD8
PCYT1A
LPCAT3
KIF13B
GPR89B
GPR89A
GM2A
PAFAH1B1
ATP2B1
CCPG1
BZW2
MYADM
SLC33A1
CDS2
ASAH1
LPGAT1
RNA metabolism
Gene expression
Regulation of gene expression
Infectious disease
Immune system

B

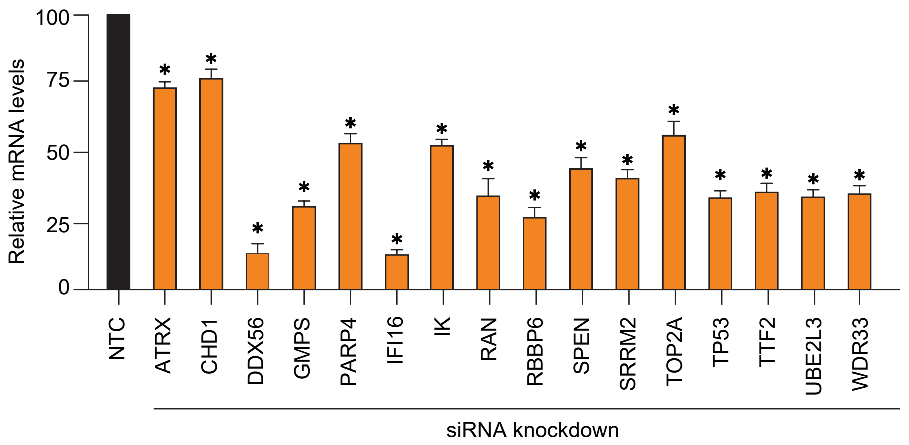
Relative mRNA levels
100
75
50
25
0
NTC
ATRX
CHD1
DDX56
GMPS
PARP4
IFI16
IK
RAN
RBBP6
SPEN
SRRM2
TOP2A
TP53
TTF2
UBE2L3
WDR33
siRNA knockdown

Figure S4.

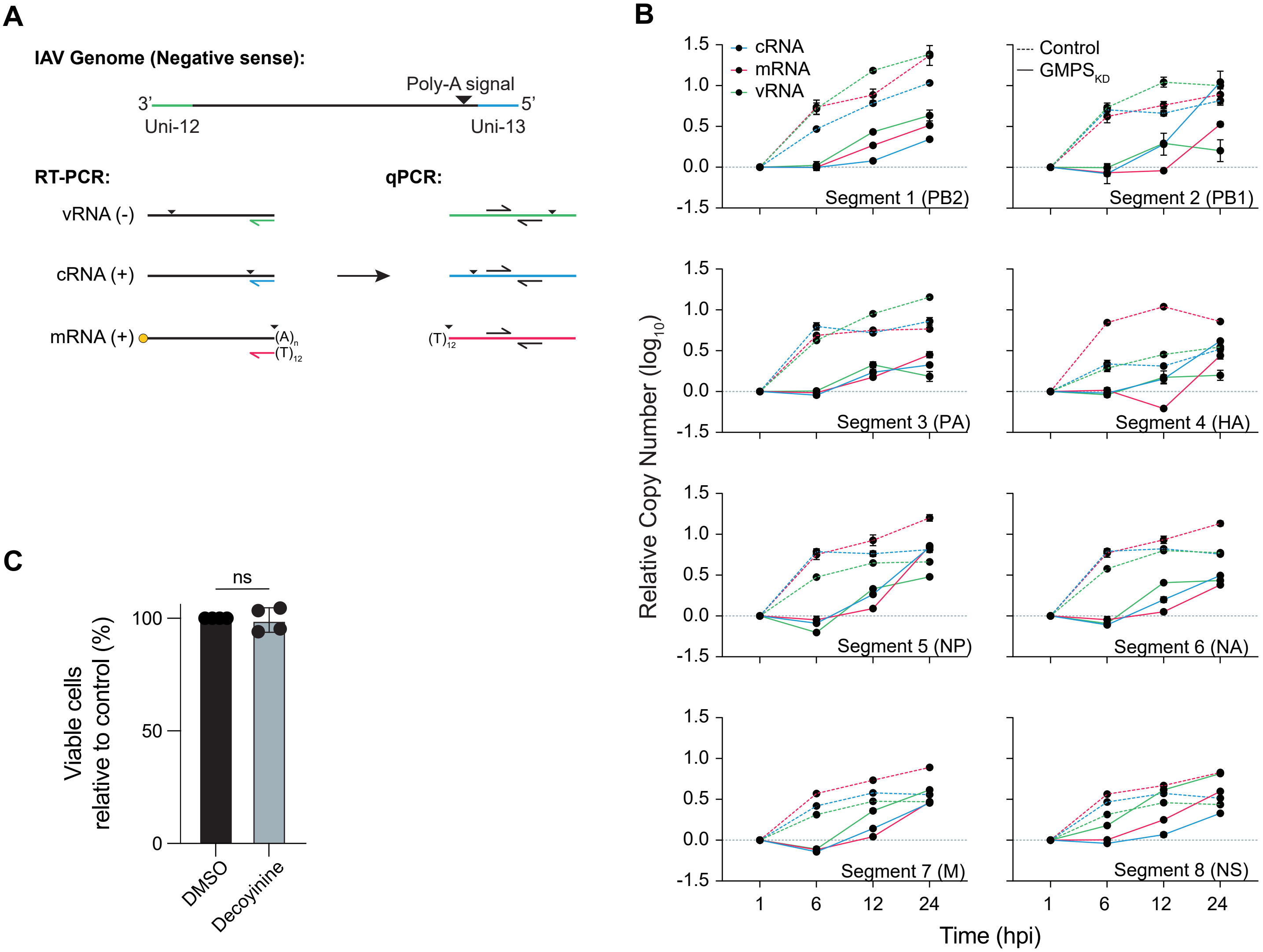
A
IAV Genome (Negative sense):
Poly-A signal
3'
5'
Uni-12
Uni-13
RT-PCR:
qPCR:
vRNA (-)
cRNA (+)
mRNA (+)
(A)n
(T)12
(T)12
B
cRNA
mRNA
vRNA
Control
GMPSKD
Segment 1 (PB2)
Segment 2 (PB1)
Segment 3 (PA)
Segment 4 (HA)
Segment 5 (NP)
Segment 6 (NA)
Segment 7 (M)
Segment 8 (NS)
Relative Copy Number (log10)
1.5
1.0
0.5
0.0
-1.5
1
6
12
24
Time (hpi)
C
ns
Viable cells relative to control (%)
100
50
0
DMSO
Decoyinine

Figure S5.

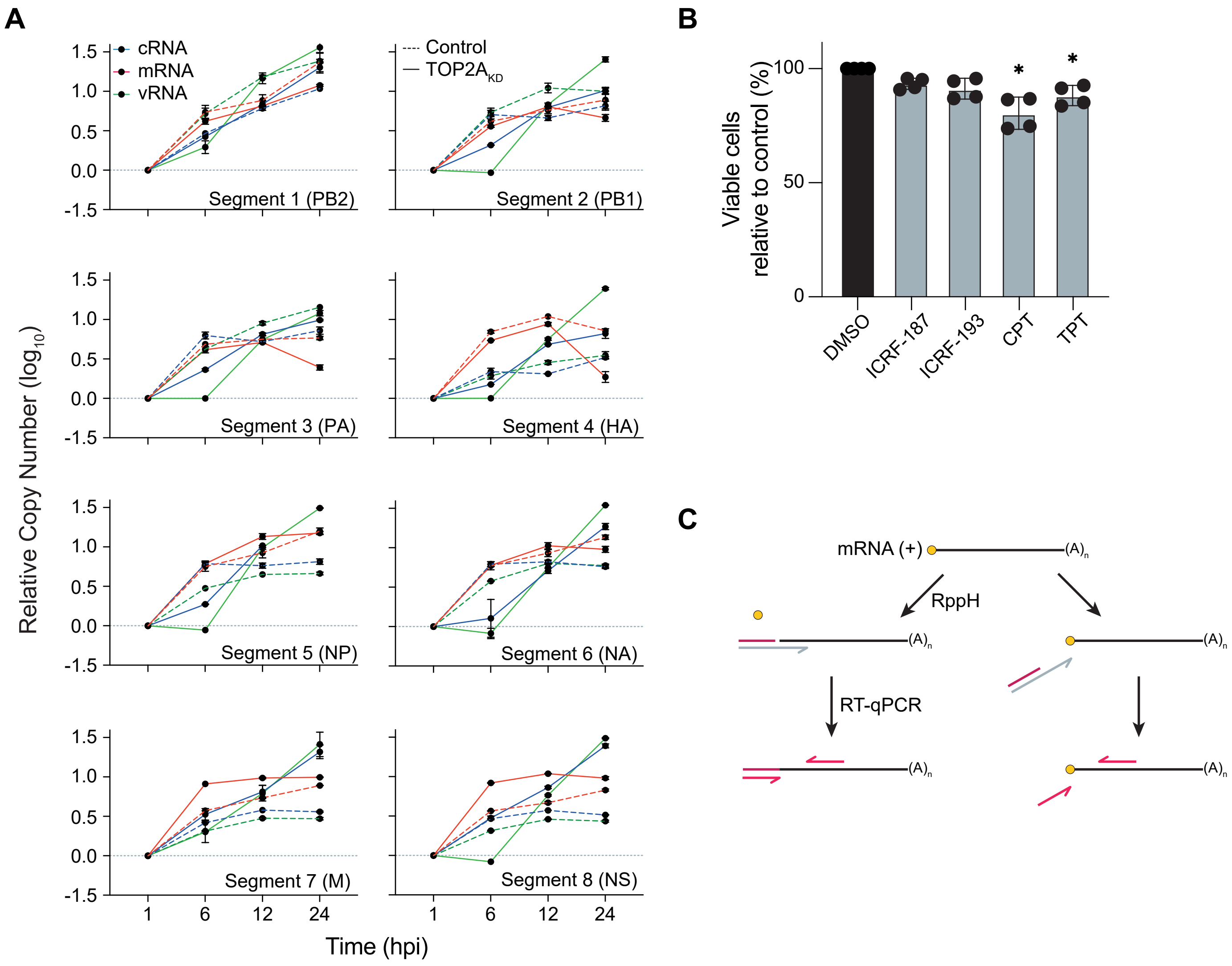
A
cRNA
mRNA
vRNA
Control
TOP2A$_{KD}$
Segment 1 (PB2)
Segment 2 (PB1)
Segment 3 (PA)
Segment 4 (HA)
Segment 5 (NP)
Segment 6 (NA)
Segment 7 (M)
Segment 8 (NS)
Relative Copy Number ($log_{10}$)
Time (hpi)
1
6
12
24
B
Viable cells relative to control (%)
DMSO
ICRF-187
ICRF-193
CPT
TPT
C
mRNA (+)
(A)n
RppH
RT-qPCR

### Figure S6.

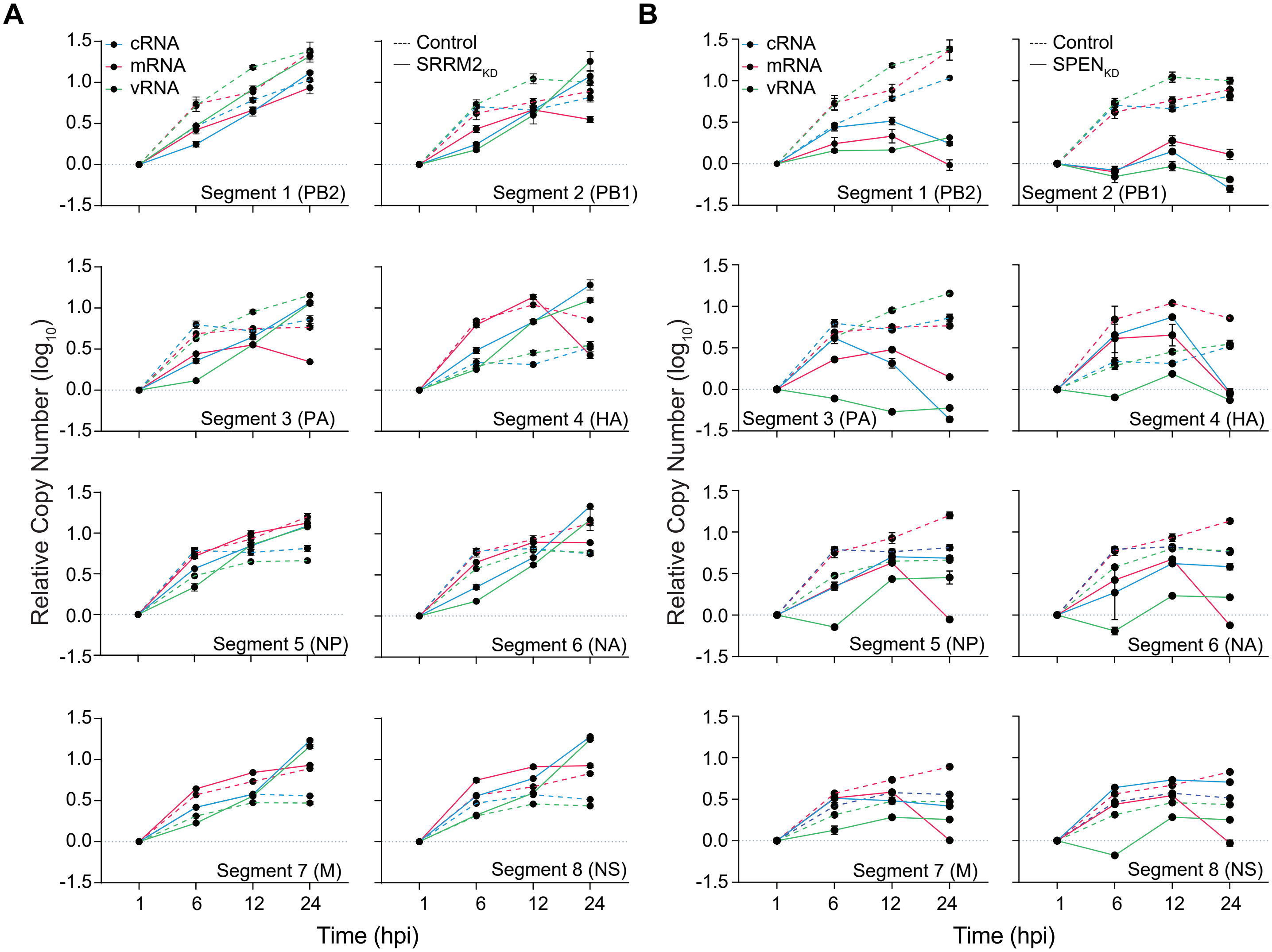
A
B
cRNA
mRNA
vRNA
Control
SRRM2KD
SPENKD
Relative Copy Number (log10)
Time (hpi)
Segment 1 (PB2)
Segment 2 (PB1)
Segment 3 (PA)
Segment 4 (HA)
Segment 5 (NP)
Segment 6 (NA)
Segment 7 (M)
Segment 8 (NS)
1.5
1.0
0.5
0.0
-1.5
1
6
12
24
